# Connexin-36-expressing Gap Junctions in VTA GABA Neurons Sustain Opiate Dependence

**DOI:** 10.1101/2020.12.18.423554

**Authors:** Geith Maal-Bared, Mandy Yee, Erika K. Harding, Martha Ghebreselassie, Michael Bergamini, Roxanne Choy, Ethan Kim, Stephanie Di Vito, Maryam Patel, Mohammadreza Amirzadeh, Taryn E. Grieder, James I. Nagy, Robert P. Bonin, Derek van der Kooy

## Abstract

Drug dependence is characterized by a switch in motivation wherein a positively reinforcing substance becomes negatively reinforcing. Ventral tegmental area (VTA) GABA neurons form a point of divergence between two double dissociable pathways responsible for these respective motivational states. Here we show that this switch from drug-naïve to opiate-dependent and withdrawn (ODW) motivation is contingent upon the gap junction-forming protein, connexin-36 (Cx36), in VTA GABA neurons. Intra-VTA infusions of the Cx36 blocker, mefloquine, in ODW rats resulted in a reversion to a drug-naïve motivational state and a loss of opiate withdrawal aversions. Consistent with these data, conditional knockout mice lacking Cx36 in GABA neurons (*GAD65-Cre;Cx36^fl(CFP)/fl(CFP)^*) were perpetually drug-naïve and never experienced opiate withdrawal aversions. Further, viral-mediated rescue of Cx36 in VTA GABA neurons was sufficient to restore their susceptibility to ODW motivation. Our findings reveal a functional role for VTA gap junctions that has eluded prevailing circuit models of addiction.

**Significance:** The motivation to seek drugs can vary depending on prior exposure. For instance, recreational and habitual drug use can stem from a desire to experience the pleasurable or relieving properties of the substance, respectively. Here we identify a subpopulation of midbrain neurons that dictate opiate-seeking motivation via expression of the gap junction protein, connexin-36. We show that connexin-36 expression increases upon opiate dependence and withdrawal. We then demonstrate that this is not merely a correlation, as pharmacological or genetic manipulations that interfere with connexin-36 function prevent the development of opiate dependence in rats and mice. Our results identify gap junctions as a critical node in the pathogenesis of opiate addiction, and a potential new target for substance use disorder pharmacotherapies.

---

The motivation to seek opiates shifts, over time and with repeated use, from a form of positive reinforcement (i.e., pleasure-seeking) to one of negative reinforcement (i.e., withdrawal avoidance; Koob and Volkow, 2010). This motivational switch is linked to a functional switch in the anatomical systems underlying drug-seeking behaviour such that the pathways responsible for opiate reinforcement are dissociable in opiate-naïve and ODW animals (Fig. 1A and 1B). The tegmental pedunculopontine nucleus (TPP) is implicated in positive reinforcement in opiate-naïve animals, but not in the negative reinforcement sought by ODW animals, while the inverse is true for mesolimbic DA. Thus, lesions of the TPP block heroin self-administration but not expression of conditioned bar pressing for heroin (Olmstead et al., 1998), and abolish morphine conditioned place preferences (CPP) in opiate-naïve animals but not ODW animals (Bechara and van der Kooy, 1992). By contrast, systemic or nucleus accumbens (NAc)-directed administrations of the broad-spectrum dopamine (DA) receptor antagonist, α-flupenthixol, abolish morphine CPP in ODW but not opiate-naïve animals. This double dissociation – wherein the TPP mediates opiate-naïve reinforcement and mesolimbic DA mediates ODW reinforcement– extends to other reinforcers such as food, ethanol, and nicotine (Bechara and van der Kooy, 1992; Laviolette et al., 2002; Nader and van der Kooy, 1997; Nader et al., 1994; Ting-A-Kee et al., 2009). Moreover, it is replicable when morphine is injected directly into the VTA, suggesting the midbrain region may act as a point of divergence for the two dissociable pathways (Nader and van der Kooy, 1997; Figs. 1*A*, 1*B*).

**Figure 1.**
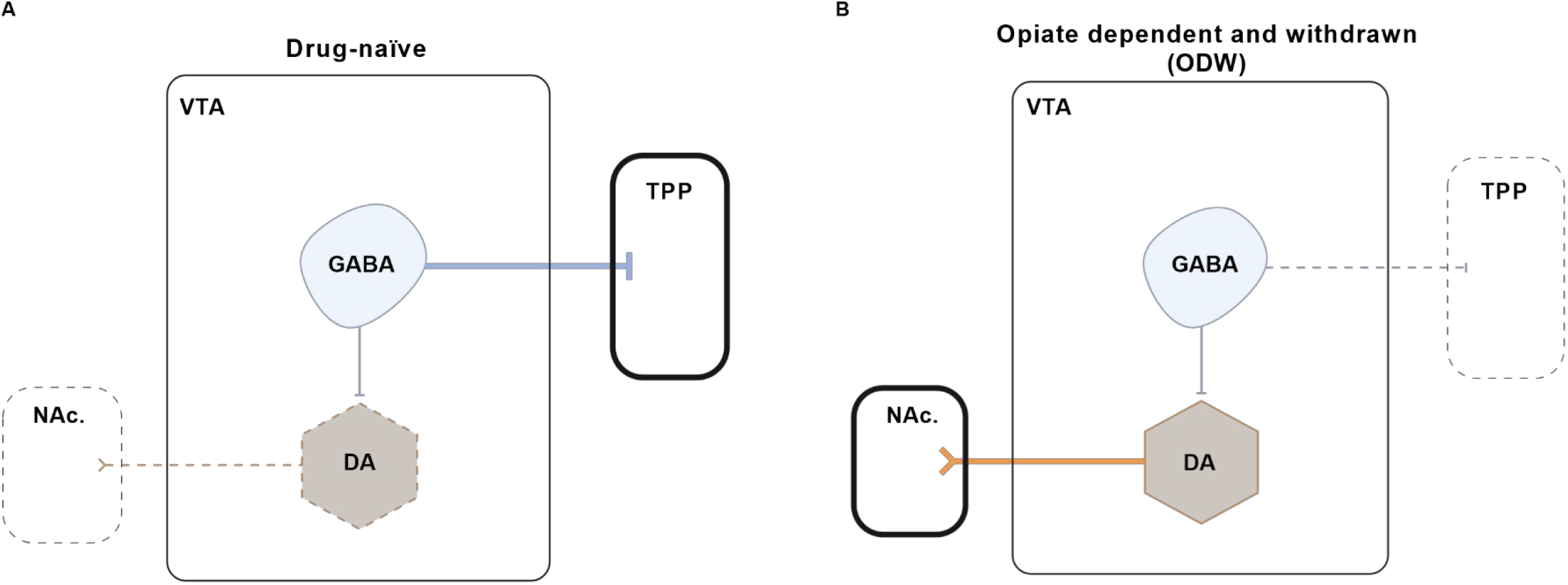
Anatomical regions implicated in opiate reinforcement in the drug-naïve and ODW motivational states. **(A and B)** Schematic diagrams depicting the double dissociation between TPP-mediated positive reinforcement for opiates in drug-naïve animals **(A)** vs. DA-mediated negative reinforcement for opiates in ODW animals **(B)**. Thick lines and borders represent structures that are functionally important for each respective motivational state, while dashed lines and borders represent structures that are not.

Several lines of evidence indicate that VTA GABA neurons may house the motivational switch between opiate positive and negative reinforcement. Single-unit extracellular recordings under normal physiological conditions (i.e., a drug-naïve state) show that VTA GABA neurons are hyperpolarized by bath application of the GABA_A_ receptor agonist, muscimol. In ODW brains, roughly half of the VTA GABA population exhibits a hyperexcitable phenotype wherein muscimol elicits depolarization, consistent with a positive shift in the GABA_A_ reversal potential and a dysregulation of chloride gradients (Laviolette et al., 2004). Most important, manipulations that impart this electrophysiological profile (e.g. intra-VTA infusions of brain-derived neurotrophic factor or blockers of the K-Cl cotransporter KCC2) result in an ODW-like behavioural phenotype in otherwise drug-naïve animals (Ting-A-Kee et al., 2013; Vargas-Perez et al., 2009). Thus, the GABA_A_ switch exhibited by VTA GABA neurons (hereafter referred to as *switched* GABA neurons) is not merely a correlate of the ODW motivational state, it is also causative. This hints that aberrations in chloride homeostasis contribute to the pathogenesis of chronic opiate use, a mechanism that seems to be implicated in other neuropathologies such as epilepsy, neuropathic pain and autism (Barmashenko et al., 2011; Coull et al., 2005; Tyzio et al., 2014). In the case of epilepsy, electrical synapses formed by gap junctions may serve as a medium by which seizure activity can spread across a network of neurons. Consistent with this, drugs that block gap junctions have anticonvulsant effects (Franco-Pérez et al., 2015; Volman et al., 2013). Some VTA GABA neurons express the neuronal gap junction protein, Cx36, and exhibit the characteristics of electrically-coupled networks (Allison et al., 2006). Whether these properties have any bearing on the motivational switch is unclear, though increased VTA GABA synchrony has been shown to increase DA burst firing (Morozova et al., 2016). On the basis that electrical synapses formed by Cx36-containing gap junctions could conceivably facilitate this synchrony and the consequent DA bursts, we hypothesized that manipulations that prevent Cx36-mediated coupling in VTA GABA neurons would interfere with the manifestation of a DA-mediated ODW motivational state. To this end, we performed a series of morphine CPP and conditioned place avoidance (CPA) assays in rats that received intra-VTA infusions of the Cx36 blocker, mefloquine, as well as conditional knockout mice lacking the Cx36 gene in GABAergic neurons (*GAD65-Cre;Cx36^fl(CFP)/fl(CFP^)*). We also performed retrograde tracing and immunohistochemistry to characterize this population of VTA GABA neurons that we demonstrate to be crucial for the development of opiate dependence.

## Materials and Methods

### Subjects

Animal use was conducted in accordance with the guidelines set forth by the University of Toronto Animal Care Committee, the Animals for Research Act in Ontario and the Canadian Council on Animal Care.

Adult male Wistar rats (250 – 400 grams) and C57BL/6 mice (20 – 35 grams) were acquired from Charles River (Montreal, Canada). *Gad2-IRES-Cre* (GAD65-Cre) and *Gad2-T2a-NLS-mCherry* (GAD65-mCherry) mice were acquired from The Jackson Laboratory (Stock No: 010802 and 023140). *Creb1^tm1Gsc^* mice (CREB hypomorphs) were obtained from Dr. Alcino Silva’s repository at The Jackson Laboratory (Stock No: 004445) via Dr. Sheena Joselyn (University of Toronto, Canada). Cx36flox(CFP) mice were acquired from the European Mutant Mouse Archive (EMMA ID: EM 02510). Cx36-EGFP mice were donated by J. Nagy (Spinal Cord Research Centre, University of Manitoba). Gad2-IRES-Cre and Cx36flox(CFP) mice were crossed to generate *GAD65-Cre;Cx36^fl(CFP)/fl(CFP)^* mice. Rats and mice were double-housed and group-housed with littermates, respectively. All animals were housed in clear Plexiglass cages containing environmental enrichment (chew toy and red-tinted plastic tunnels) in the Division of Comparative Medicine at the University of Toronto. Ambient temperature was maintained at 22°C for both the rat and mouse colony rooms. All subjects were kept on a 12-hour light/dark cycle (lights on at 7 AM, lights off at 7 PM). All subjects had *ad libitum* access to food and water.

### Drugs

The drugs used in our experiments were: morphine sulfate (Almat Pharmachem), diacetylmorphine hydrochloride (heroin; Almat Pharmachem), mefloquine hydrochloride (Sigma-Aldrich), cis-(Z)-flupenthixol dihydrochloride (α-flu; Sigma-Aldrich), muscimol hydrobromide (Sigma-Aldrich), furosemide (Sigma-Aldrich), naloxone hydrochloride dihydrate (Sigma-Aldrich), lithium chloride (LiCl; Sigma-Aldrich), lidocaine hydrochloride (Sigma-Aldrich), N-Methyl-D-aspartic acid (NMDA; Sigma-Aldrich) and sodium pentobarbital (Ceva Santé Animale). All doses were selected based on previous work (Bissiere et al., 2011; Dockstader et al., 2001; Mucha et al., 1982; Vargas-Perez et al., 2014).

For intracranial injections (I.C.), rats were surgically implanted with bilateral cannulas (Plastics One) targeting the VTA or the TPP. During the microinfusions, each hemisphere received 0.3 uL of drug or vehicle over the span of one minute. Injector tips were left in place for an additional minute to ensure drug diffusion into the target region. Injectors were removed over the span of one minute to minimize solution displacement. The drugs administered via intracranial injections were mefloquine, furosemide, lidocaine, and NMDA. Mefloquine (100 mM) was dissolved in 0.1 M sterile PBS containing 1% DMSO. Furosemide (1 mM) and lidocaine (4%) were dissolved in PBS (pH = 7.4). We also used a cocktail of mefloquine and furosemide to assess the relationship between Cx36 and KCC2 in the motivational switch. The cocktail contained 100 mM mefloquine and 1 mM furosemide dissolved in 0.1 M PBS containing 1% DMSO. Control groups received PBS or PBS containing 1% DMSO – whichever vehicle corresponded with the experimental group’s treatment.

Mice used for retrograde tracing experiments received cholera toxin b subunit (CTB) conjugated with Alexa Fluor 488 and/or Alexa Fluor 647 (CTB-488 and CTB-647; ThermoFisher, C34775 and C34778).

All systemic drug administrations in both rats and mice had a volume of 0.25 mL. The drugs injected systemically were morphine (10 mg/kg, I.P. in rats; 5 mg/kg, I.P. in mice), heroin (0.5 mg/kg, S.C. in rats), α-flu (0.8 mg/kg, I.P. in rats; 0.5 mg/kg, I.P. in mice), naloxone (0.1 mg/kg S.C. in mice) and lithium chloride (5 mg/kg, I.P. in rats and mice). All systemically administered drugs were dissolved in PBS (pH = 7.4). Control groups received systemic PBS.

### Opiate dependence induction

#### Rat Dependence Induction

In rats, opiate dependence was induced through systemic injections of heroin (0.5 mg/kg; S.C.) Rats received one injection per day for five days prior to the first day of conditioning. During the eight days of conditioning, dependence was maintained through administrations of the same dose of heroin roughly 3 hours after conditioning. The next conditioning session was conducted 21 hours following heroin administration – a timepoint of spontaneous opiate withdrawal. Rats received a total of thirteen heroin injections – the last of which occurred on the eighth and final day of conditioning. Rats were tested at least 48 hours following the last heroin administration.

#### Mouse Dependence Induction

In mice, opiate dependence was induced via subcutaneous osmotic pumps (Model 1007D, Alzet Osmotic Pumps) filled with morphine solution (60 mg/mL)(Dockstader et al., 2001). Controls received pumps containing PBS. The pumps delivered 0.5 μl of solution per hour. For experiments assaying morphine place preferences, the first conditioning session began 24 hours following pump implantation. Withdrawal was precipitated at the beginning of mCPP training sessions with naloxone (0.1 mg/kg; S.C.) after which morphine or saline were administered. Pumps were explanted after the eighth and final conditioning session. Mice were tested at least 48 hours following pump removal.

For experiments assaying opiate withdrawal aversions, mice received the pumps and were left unperturbed in their homecage for 5-7 days, after which the pumps were explanted. Mice were conditioned 12 – 16 hours following pump removal – a timepoint of spontaneous opiate withdrawal at which we have observed mice to exhibit CPA. Mice were tested at least 48 hours following the single conditioning session. This protocol elicits withdrawal aversions that are qualitatively similar to that of our heroin pre-exposure protocol used in rats(Dockstader et al., 2001).

### Surgical procedures

#### Rat surgeries

For recurring drug administrations directed to the VTA or TPP, rats were surgically cannulated in a stereotaxic surgery setup at least one week prior to dependence induction. Anesthesia was induced and maintained with 4% and 2-3% isoflurane, respectively. Rats received the analgesic, ketoprofen (5 mg/kg; S.C.), during surgery and twice daily for two days post-op. The scalp was shaved and cleaned with ethanol and betadine. Once placed in the stereotaxic frame, the skull was exposed and burr holes were drilled above the target region. To target the VTA, 22-gauge stainless steel guide cannulae (Plastics One) were lowered at a 10° angle into the following coordinates relative to bregma: AP: −5.3 mm; ML: ± 2.3 mm and DV: −8.0 mm. TPP cannulations were at a 10° angle at the following coordinates: AP: −7.6 mm; ML: ± 3.1 mm and DV: −6.6 mm (Paxinos and Watson, 2006). For TPP inactivation, we performed TPP cannulation surgeries and administered I.C. lidocaine prior to each conditioning session.

All rats had a post-operative recovery period of at least one week following craniotomy.

#### Mouse surgeries

For mouse surgeries, isoflurane was used as an anesthetic (3% induction, 2% maintenance) and meloxicam (5 mg/kg; S.C.) was used as an analgesic. For dependence induction, osmotic pumps containing morphine (6 mg/pump) or PBS were planted subcutaneously under aseptic conditions. Viral and tracer delivery in mice were performed by delivering 0.3 μL solution bilaterally via 1 μL Hamilton Syringes. The protocol for stereotaxic surgery in mice was similar to the one used for rats. For viral delivery into the VTA, our coordinates were identical to ones used in our previous work (Vargas-Perez et al., 2014; AP: −3.00 mm, ML: ±0.5 mm, DV: −4.2 mm from bregma). These coordinates were used to deliver Cre-dependent AAVs carrying Cx36 (ssAAV9-Ef1a-DIO-Cx36) and EGFP (ssAAV9-Ef1a-EGFP-WPREs) at 5E+13 GC/mL in 0.1 mL. AAVs were obtained from Packgene. Coordinates for retrograde tracing were AP: −4.4 mm, ML: ± 1.25 mm, DV: −3.75 mm for the TPP and AP: +1.7 mm, ML: ± 1.00 mm, DV: −4.00 mm for the NAc.

#### Place conditioning in rats

Rats were left to recover for at least one week following craniotomy before any injections (e.g. dependence induction) were administered and any conditioning was conducted (Fig. 3a). To minimize confounds of novelty, rats were habituated for 20 minutes in a neutral grey chamber (41 cm x 41 cm x 38 cm) that resembles our place conditioning boxes (day-1). To further reduce any novelty effects elicited by our conditioning paradigm, rats underwent a mock infusion session – in which I.C. saline was administered – on the following day (day 0). Place conditioning was conducted on the following eight days (day 1 – day 8). We used an unbiased procedure as rats do not exhibit baseline preferences in our paradigm (Dockstader et al., 2001; Mucha et al., 1982). Each conditioning session was 40 minutes long. In experiments assaying morphine place preferences, drug-place pairings and first day of drug exposure were counterbalanced across all groups. In experiments assaying opiate withdrawal aversions, withdrawal-place pairings were counterbalanced. Our place conditioning apparatus consists of two boxes (41 cm × 41 cm × 38 cm) that are distinguishable visually (black vs. white), tactilely (solid surface vs. grid surface) and olfactorily (~0.3 mL glacial acetic acid vs. unscented).

For experiments assaying morphine place preferences, the conditioning period lasted eight days. Morphine and PBS were administered on alternating days such that rats received four drug-place and four vehicle place pairings. Conditioning sessions always occurred 21 hours following the last heroin or vehicle injections. For experiments assaying opiate withdrawal aversions, the conditioning period lasted eight days but rats were only conditioned four times. Rats were placed in one of the two compartments on the first, third, fifth and seventh days – always at a timepoint coinciding with spontaneous opiate withdrawal (21 hours following the last heroin or vehicle injection). In so doing, rats received four withdrawal-place pairings interspersed with four days in the homecage. Opiate dependence was induced prior to the pre-conditioning habituation sessions and maintained until the last day of conditioning. For experiments with I.P. pretreatments, α-flu or PBS were administered 2.5 hours before conditioning. For I.C. pretreatments, mefloquine, furosemide, the cocktail of mefloquine and furosemide, or DMSO were administered 40 minutes prior to conditioning, given that the non-specific effects of mefloquine diminish at the 40-minute timepoint (Allison et al., 2011). The first day of conditioning was the sixth day of heroin administration for opiate-dependent rats. For all behavioural experiments, we conducted testing sessions at least 48 hours following the final day of conditioning to minimize any residual effects of the drugs administered during the conditioning period. Testing sessions were 10 minutes long.

For morphine CPP experiments, difference scores were calculated by subtracting time spent in the vehicle-paired compartment from the time spent in the morphine-paired compartment. For withdrawal CPA experiments, we subtracted time spent in the withdrawal-paired compartment from the time spent in the neutral compartment.

Noldus Ethovision XT was used for all rat CPP and CPA data acquisition.

#### Somatic signs of opiate-withdrawal

Abstinence scores were generated by recording the number of somatic withdrawal signs (e.g., wet dog shakes, gnawing, scratching, spasms, and audible vocalizations) 1, 16 and 21 hours following the last heroin injection. Observation periods were 30 minutes long. The absolute number of somatic withdrawal signs observed in the DMSO-treated naïve rats was used as a baseline to which the other groups were compared.

#### Place conditioning in mice

An unbiased procedure was used as our mice do not show a baseline preference for either compartment in our paradigm (Dockstader et al., 2001). For morphine CPP experiments, an eight-day conditioning period began the day after osmotic pumps (containing morphine or saline) were implanted. For morphine CPP experiments, drug-place pairings and first day of drug exposure were counterbalanced. For withdrawal CPA experiments, withdrawal-place pairings were counterbalanced. Mice with morphine pumps (i.e. ODW mice) received naloxone injections (0.1 mg/kg SC) for withdrawal precipitation 5 minutes prior to morphine (5 mg/kg IP) or saline administration. Mice were placed in the conditioning apparatus immediately following the administration of morphine or saline. Morphine place conditioning sessions in mice were 15 minutes long.

For withdrawal CPA experiments, no naloxone was administered to precipitate opiate withdrawal. Instead, mice were left in their homecage for 6-7 days following osmotic pump planting, after which pumps were removed to induce spontaneous opiate withdrawal. Mice underwent a single, 50-minute long conditioning session 12-16 hours following pump removal. Both morphine CPP and withdrawal CPA testing sessions were 10 minutes long.

The mouse place conditioning apparatus (Place Conditioning Chamber, Med Associates Inc.) consisted of two different environments measuring 15 cm × 15 cm × 15 cm. Similar to the rat conditioning apparatus, the two compartments were visually (black vs. white) and tactilely (metal rod floor vs. wire mesh floor) dissociable. Each compartment was cleaned immediately after use.

Mice used for CPP experiments were *GAD65-Cre;Cx36^fl(CFP)/fl(CFP)^* and littermate controls lacking the GAD65 promoter-driven Cre recombinase (i.e. *Cx36^fl(CFP)/fl(CFP)^* mice).

The Med-PC IV software suite was used for all mouse CPP and CPA data acquisition

#### Histology

Subjects were deeply anesthetized with sodium pentobarbital (54 mg/kg; I.P.) and transcardially perfused with 0.9% saline followed by cold 4% paraformaldehyde (PFA; Sigma Aldrich, P6148). The brains were extracted and post-fixed overnight in 4% PFA. Brains were then cryoprotected in 30% sucrose-containing PBS. Brains were sectioned coronally using a cryostat. Rat brains were sectioned at 30 μm thickness, mounted on gelatin-coated slides and stained with cresyl violet for cannula placement verifications via light microscopy. Mouse brains were sectioned at a 10 μm thickness and mounted on charged slides for injection track verification via light microscopy or immunostaining and confocal microscopy.

#### Immunohistochemistry and microscopy

Brain slices were washed with PBS (pH = 7.4) for two 5-minute intervals followed by a 10-minute interval. Sections were then blocked with 5% normal goat serum (NGS) containing 2% bovine serum albumin (BSA) and 0.2% Triton-X 100 for a duration of 1 hour at room temperature. The slides were then washed with PBS thrice and incubated with the appropriate primary antibody overnight at 4°C. Unless otherwise stated, all primary antibodies were diluted in PBS containing 1% NGS and 1% BSA. A wide variety of primary antibodies was used in our experiments: rabbit polyclonal anti-TH (1/1000; Abcam, ab112), mouse monoclonal anti-GFP (1/500; Abcam, ab1218), rabbit polyclonal anti-GFP (1/3250; Thermofisher, PA1-980A), rabbit monoclonal anti-GFP (1/200; Thermofisher, G10362) mouse monoclonal anti-pCREB (1/250; Millipore Sigma, 05-807), and rabbit polyclonal anti-Cx36 (1/250; Thermofisher, 36-4600).

On the following day, slides were washed with PBS then incubated with the secondary antibody diluted in PBS containing 1% NGS and 1% BSA for a duration of 1 hour at room temperature. Where appropriate, we used a goat anti-mouse IgG secondary antibody conjugated with Alexa Fluor 488 (1/600; Thermofisher, A28175) and/or a rabbit anti-goat IgG secondary antibody conjugated with Alexa Fluor 568 (1/600; Thermofisher, A-11079).

Following incubation with the secondary antibody, sections were washed with PBS then counterstained with Hoechst (1/1000) for 7-10 minutes at room temperature. For experiments investigating two antigens, we combined the primary antibodies into one solution and on the following day, combined the two secondary antibodies as well. After nuclear staining and another set of PBS washes, Fluoro-Gel mounting media was applied. Slides were coverslipped and sealed with CoverGrip (Biotium). Slides were stored in a cold room (4°C) for short-term use or a freezer (−20°C) for durations exceeding two weeks.

Fluorescent specimens were imaged with either a Zeiss inverted fluorescence microscope or a confocal microscope. Images were captured with Fluoview software and analyzed using ImageJ. ImageJ was also used for fluorescence intensity analysis. Corrected total cell fluorescence (CTCF) was measured using the following equation:

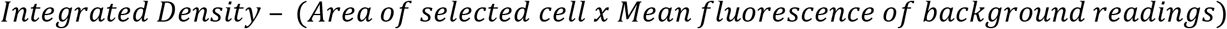

Three background fluorescence readings were obtained for each cell by selecting three regions adjacent to the region of interest (ROI).

#### Statistical analyses

Results were analyzed using a student’s *t* test or analysis of variance (ANOVA). Significance tests were displayed such that *p<0.05, **p<0.01, ***p<0.001, ****p<0.0001 as determined by student’s *t* test, one-way and two-way analysis of variance (ANOVA), or Pearson’s correlation. Data are represented as means ± standard error (SEM). Statistical analyses were performed using Graphpad Prism.

## Results

### Cx36 is upregulated in the ODW motivational state

We first sought to determine whether drug dependence has an impact on the regulation of Cx36 expression. We performed immunofluorescence labelling in drug-naïve vs. ODW mice in which enhanced green fluorescent protein serving as a reporter for Cx36 expression (Cx36-EGFP mice). Less than 1% of TH+ cells expressed Cx36-EGFP in both drug-naïve and ODW mice, corroborating previous reports that the gap junction protein is expressed primarily in GABAergic neurons (Deans et al., 2001; Galarreta and Hestrin, 2001; Fig. 2*A*). By contrast, we observed an-almost two-fold increase in the numbers of EGFP+ neurons in ODW VTA (30% ± 2%) compared to drug-naïve VTA (15% ± 1%; Figs. 2*B*, *C*). Cx36 upregulation occurred throughout the rostrocaudal axis of the VTA with a trend towards greater upregulation caudally (Figs. 3*A-D*). Fluorescence intensity in single VTA neurons also was significantly higher in ODW Cx36-EGFP mice, suggesting that individual VTA neurons express higher levels of Cx36 (Figs. 4*A*-*C*).

**Figure 2.**
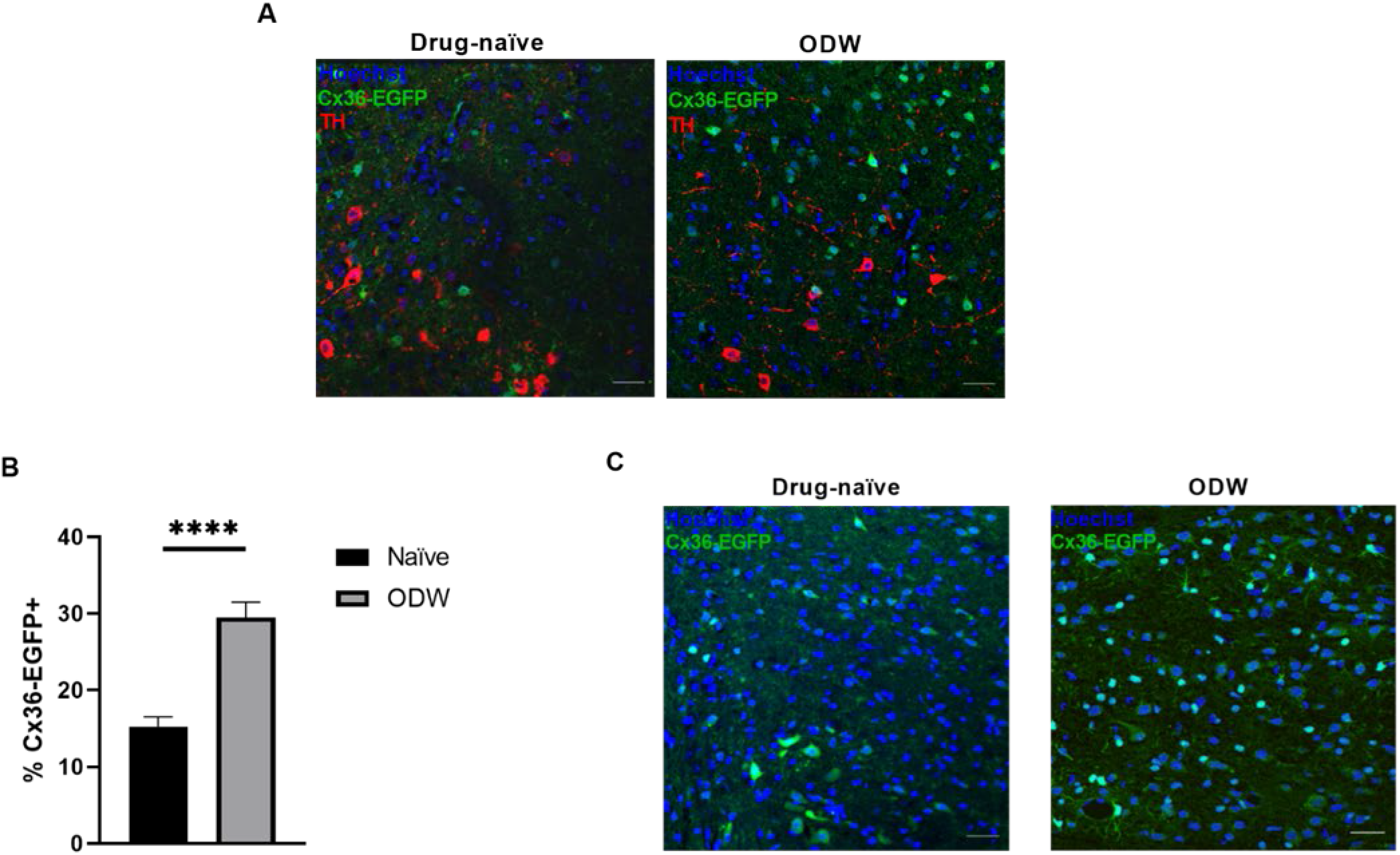
Cx36 is upregulated in GABA neurons of the VTA in ODW mice. **(A)** Representative immunofluorescence of VTA cells (Hoechst: blue) from drug-naïve and ODW Cx36-EGFP mice following GFP (Cx36-EGFP: green) and TH (red) staining. Less than 1% of TH+ neurons expressed Cx36-EGFP irrespective of motivational state (i.e. drug-naïve or ODW). **(B)** Summary data showing proportion of EGFP+ VTA cells in drug-naïve and ODW Cx36-EGFP mice. A two-fold increase in Cx36-EGFP staining was observed in ODW Cx36-EGFP mice (*n*=4 mice, 17184 VTA cells) compared to drug-naïve Cx36-EGFP mice (*n*=4 mice, 25848 VTA cells; Two-tailed *t*(102)=6.197, ****: *p*<0.0001). Data are represented as mean ± SEM. **(C)** Representative immunofluorescence of drug-naïve and ODW VTA cells (Hoechst: blue) expressing Cx36-EGFP (EGFP: green). Scale bars: 30 μm.

**Figure 3.**
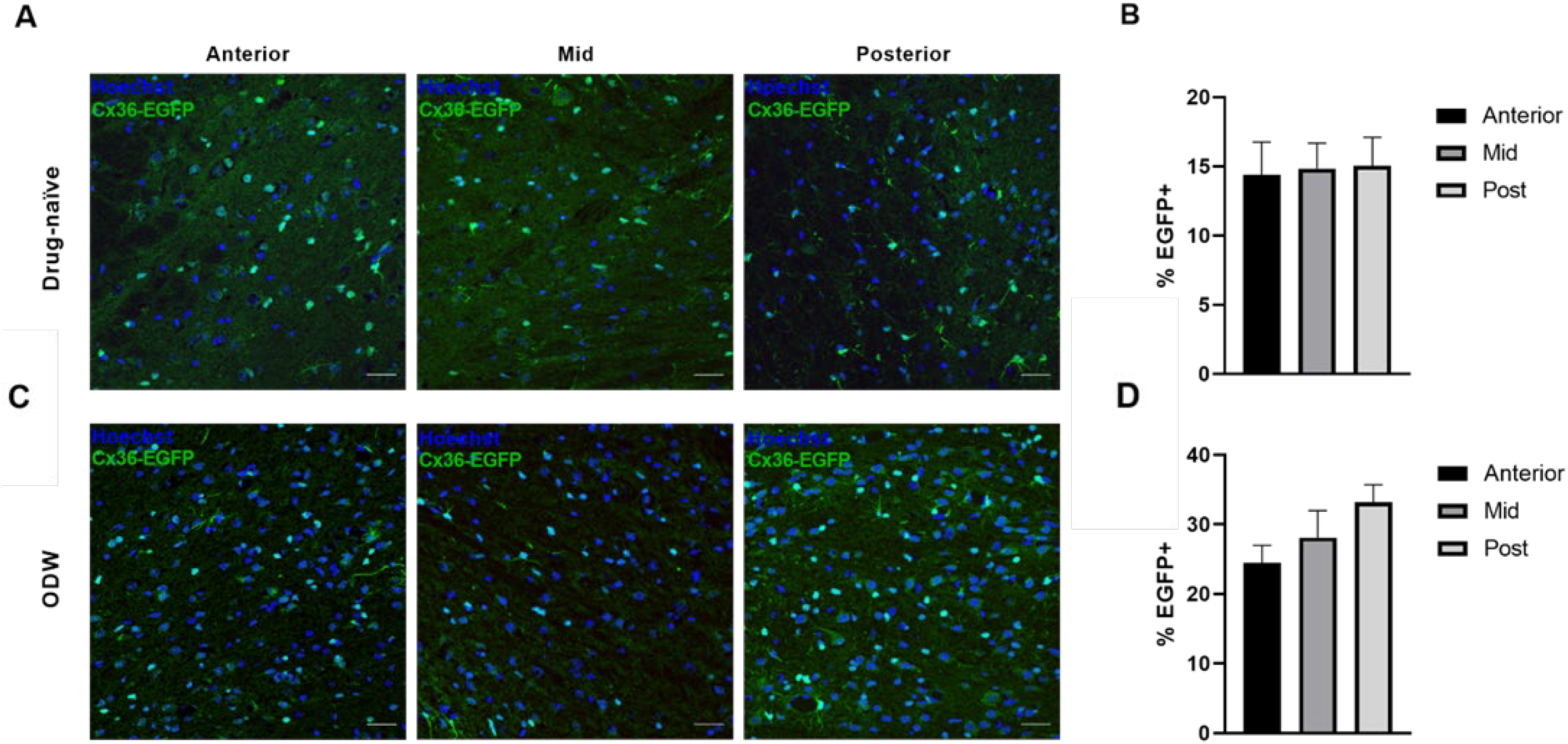
Cx36 is upregulated throughout the rostrocaudal axis of the VTA in ODW mice. **(A)** Representative immunofluorescence of anterior, mid and posterior VTA cells (Hoechst: blue) from drug-naïve Cx36-EGFP mice following GFP staining (EGFP: green). **(B)** Summary data showing the percentage of anterior, mid, and posterior VTA cells that express Cx36 (i.e. EGFP+) in drug-naïve Cx36-EGFP mice. There was no significant difference in the regional expression along the rostrocaudal axis (*F*_(2,45)_=0.02168, p=0.9786). **(C)** Representative images of immunofluorescence of anterior, mid and posterior VTA cells (Hoechst: blue) from ODW Cx36-EGFP mice following GFP staining (EGFP: green). **(D)** Summary data showing the percentage of anterior, mid, and posterior VTA cells that express Cx36 (i.e. EGFP+) in ODW Cx36-EGFP mice. Although there is a trend towards increased expression caudally, this effect was not statistically significant (*F*_(2,44)_=1.429, p=0.2505). Data are represented as mean ± SEM. Scale bars are 30 μm.

**Figure 4.**
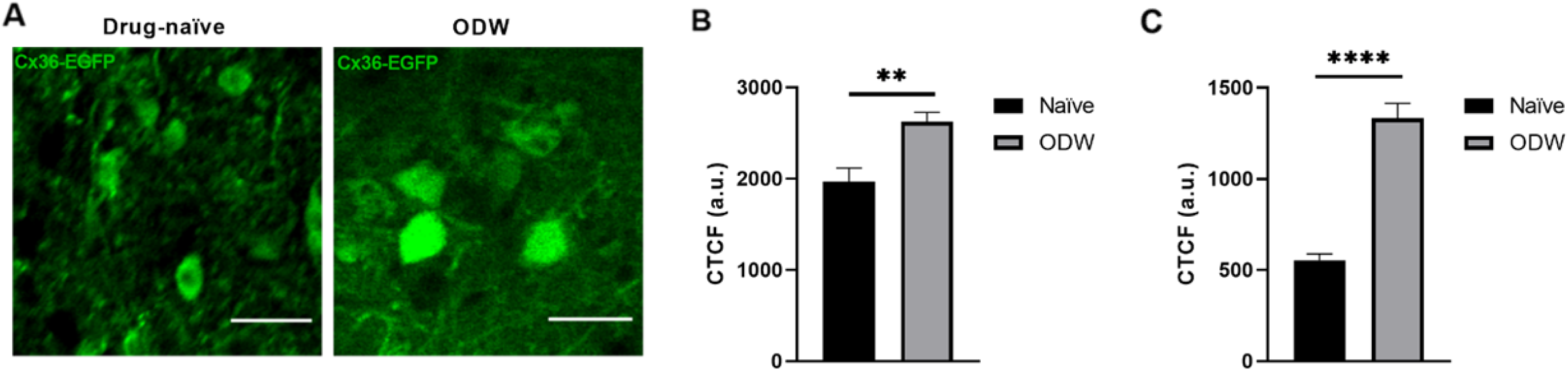
Cx36 is upregulated within individual VTA GABA neurons in ODW mice. **(A)** Representative immunofluorescence depicting the heightened intensity of endogenous Cx36-EGFP fluorescence in ODW mice compared to drug-naïve ones. **(B)** Corrected total cell fluorescence (CTCF) of native endogenous EGFP expression in VTA cells of drug-naïve and ODW Cx36-EGFP mice. CTCF values are represented in arbitrary units (a.u.) Fluorescence intensity was significantly higher in ODW VTA than drug-naïve VTA (two-tailed t-test: t_(293)_ = 3.303, p=0.0011, n = 295 VTA cells, 77 cells in 2 drug-naïve mice and 218 cells in 3 ODW mice). **(C)** CTCF of GFP-stained VTA cells of drug-naïve and ODW Cx36-EGFP mice. CTCF values are represented in arbitrary units (a.u.) Fluorescence intensity was significantly higher in ODW VTA than drug-naïve VTA (two-tailed t-test: t_(138)_=8.861, p<0.0001, n=140 VTA cells, 70 cells in 2 drug-naïve mice and 70 cells in 2 ODW mice). Data are represented as mean ± SEM. ** denotes p<0.01. **** denotes p<0.0001. Scale bars are 30 μm.

### GABA_A_ switch occurs in a majority of TPP-bound VTA neurons

We crossed a *Gad65-IRES-Cre* (GAD65-Cre) mouse line with *Cx36flox(CFP)* mice thereby generating *GAD65-Cre;Cx36^fl(CFP)/fl(CFP)^* mice, which lack Cx36 in GABAergic neurons (Taniguchi et al., 2011; Wellershaus et al., 2008). To confirm that the gene was successfully knocked out, we immunolabeled the CFP excision reporter (CFPER) and found that an average of 12% ± 1% of total VTA cells were CFPER+ in *GAD65-Cre;Cx36^fl(CFP)/fl(CFP)^* mice. Given that GABA neurons constitute 20-30% of VTA neurons, this suggests that Cx36 was originally expressed but now excised in roughly half of the VTA GABA population (Margolis et al., 2012; Nair-Roberts et al., 2008; Swanson, 1982). By contrast, littermate controls lacked CFPER expression, confirming that the Cx36 excision had occurred only in the experimental group (Figs. 5*A*, *B*).

**Figure 5.**
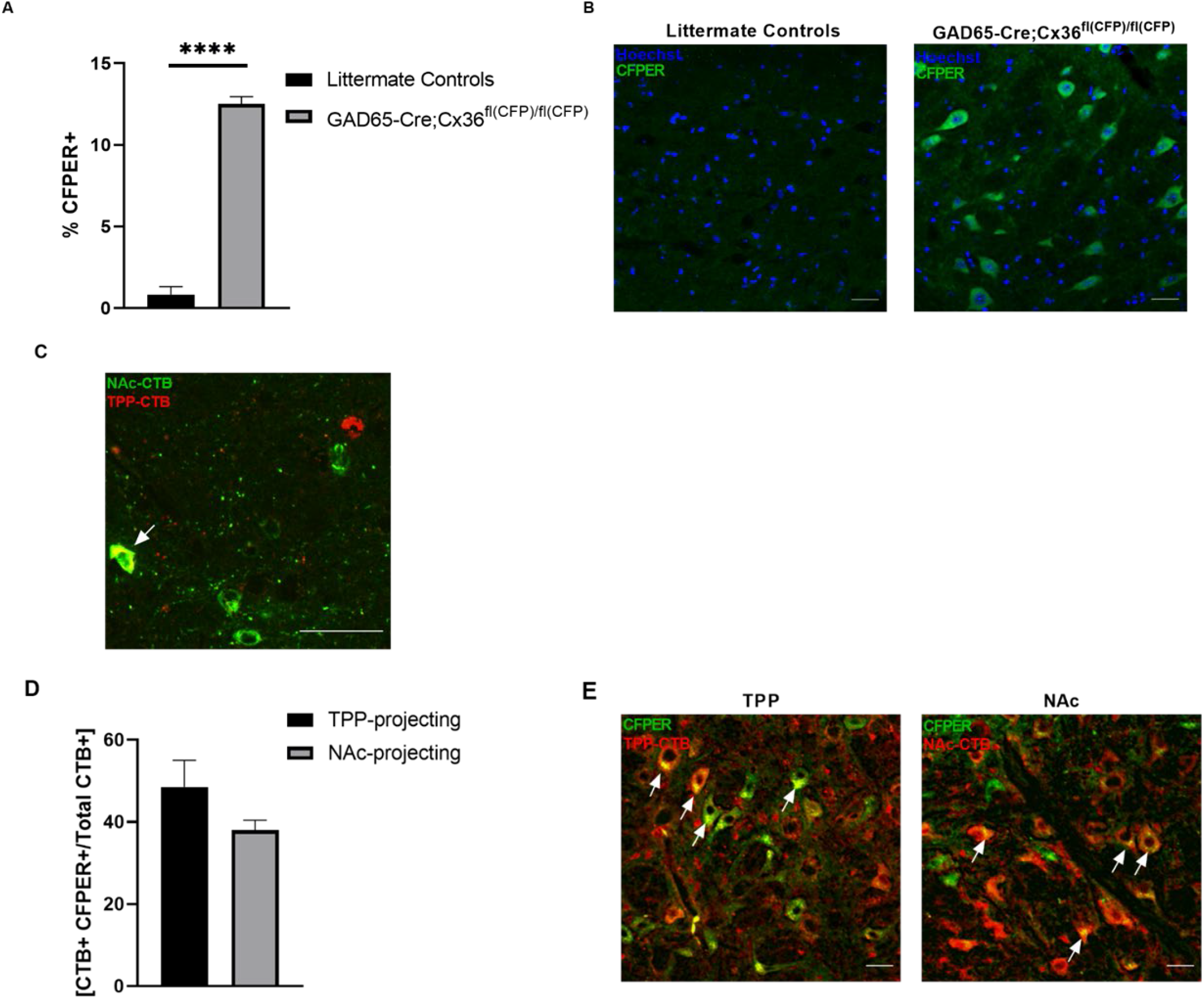
Cx36 excision reporter (CFPER) is expressed in TPP- and NAc-bound VTA neurons of *GAD65-Cre;Cx36fl(CFP)/fl(CFP)* mice. **(A)** Summary data showing proportion of VTA cells that express the Cx36 excision reporter (CFPER) in *GAD65-Cre;Cx36^fl(CFP)/fl(CFP)^* and littermate controls. CFPER expression is significantly higher in *GAD65-Cre;Cx36^fl(CFP)/fl(CFP)^* mice (*n*=15 mice, 85878 VTA cells) compared to littermate controls (*n*=6 mice, 11619 VTA cells; Two-tailed *t*_(382)_=7.480, ****: *p*<0.0001). **(B)** Representative immunofluorescence of CFPER (green) in VTA cells (blue) of *GAD65-Cre;Cx36^fl(CFP)/fl(CFP)^* mice and littermate controls. **(C)** Representative immunofluorescence depicting the VTA in wild-type mice that received CTB-488 (green) and CTB-647 (red) into the NAc. and TPP, respectively. Only about 1% of VTA cells stained co-stained for both tracers (n=4 mice). Arrow points to a double-labeled cell. **(D and E)** Immunohistochemistry of VTA cells in *GAD65-Cre;Cx36^fl(CFP)/fl(CFP)^* mice following CTB retrograde tracing from the TPP (*n=*6 mice, 48672 VTA cells) or NAc (*n=*5 mice, 33101 VTA cells). **(D)** Proportion of TPP- and NAc-projecting (CTB+) VTA cells that express CFPER. **(E)** Representative immunofluorescence of VTA cells (blue) expressing CFPER (green) and CTB (red). Arrows: representative double-labeled (yellow) cells with punctate and cytoplasmic staining.

To explore whether these CFPER+ VTA neurons had distinct projection patterns, we performed a series of retrograde tracing experiments in which cholera toxin B subunit (CTB) was injected into the TPP or NAc of *GAD65-Cre;Cx36^fl(CFP)/fl(CFP)^*mice. The negligible number of double labelled single cells that simultaneously project to both structures (Fig. 5*C*) permitted tracing each structure in separate cohorts of mice. We found that 48% ± 6% of TPP-projecting cells were CFPER+, while about 38% ± 2% of the NAc-projecting cells were CFPER+ (Figs. 5*D*, *E*).

The phosphorylated form of CREB (pCREB) is a marker for switched VTA GABA neurons (i.e. neurons depolarized rather than hyperpolarized by the GABA_A_ agonist muscimol) in ODW rats (Laviolette et al., 2004). It also may be functionally important for the motivational switch from positive to negative reinforcement based on the observation that ODW CREB homozygous hypomorphs do not exhibit morphine CPP (Figs. 6*A*, *B*). To investigate the projection patterns of switched VTA GABA neurons, we performed retrograde tracing in pCREB-stained VTA from GAD65-mCherry mice (Peron et al., 2015). First, we demonstrated that – as is the case in rats – VTA pCREB is expressed in ODW but not in drug-naïve mice (Laviolette et al., 2004). Specifically, 14% ± 1% of VTA cells were pCREB+ in ODW mice (Figs. 6*C*, *D*). Of the pCREB+ VTA neurons, 46% ± 3% were mCherry+, indicating that pCREB is expressed in GAD65-cells. Although the caudal VTA had the largest proportion of pCREB+ cells, the differences in expression along the rostrocaudal axis were not significant (Fig. 6*E*). Due to the lack of pCREB expression in drug-naïve mice, all retrograde tracing of projections of pCREB+ VTA cells was performed in ODW mice. Retrograde tracing from the TPP and NAc revealed that the vast majority (84% ± 4%) of TPP-projecting VTA cells expressed pCREB, whereas roughly half (55% ± 6%) of the NAc-projecting VTA cells were pCREB+ (Figs. 6*F*, *H*). We also observed that 56% ± 4% of the pCREB+ population was TPP-projecting and 23% ± 4% was NAc-projecting (Figs. 6*G*, *H*). It is important to note that the TPP-projecting neurons are likely GABAergic, as VTA DA neurons do not have descending projections (Swanson, 1982). Given the causal link between the GABA_A_ switch (marked by pCREB) and the motivational switch from drug-naïve to ODW (Laviolette and van der Kooy, 2001; Laviolette et al., 2004), the expression of pCREB in most TPP-bound VTA GABA neurons hints at a functional role for this projection.

**Figure 6.**
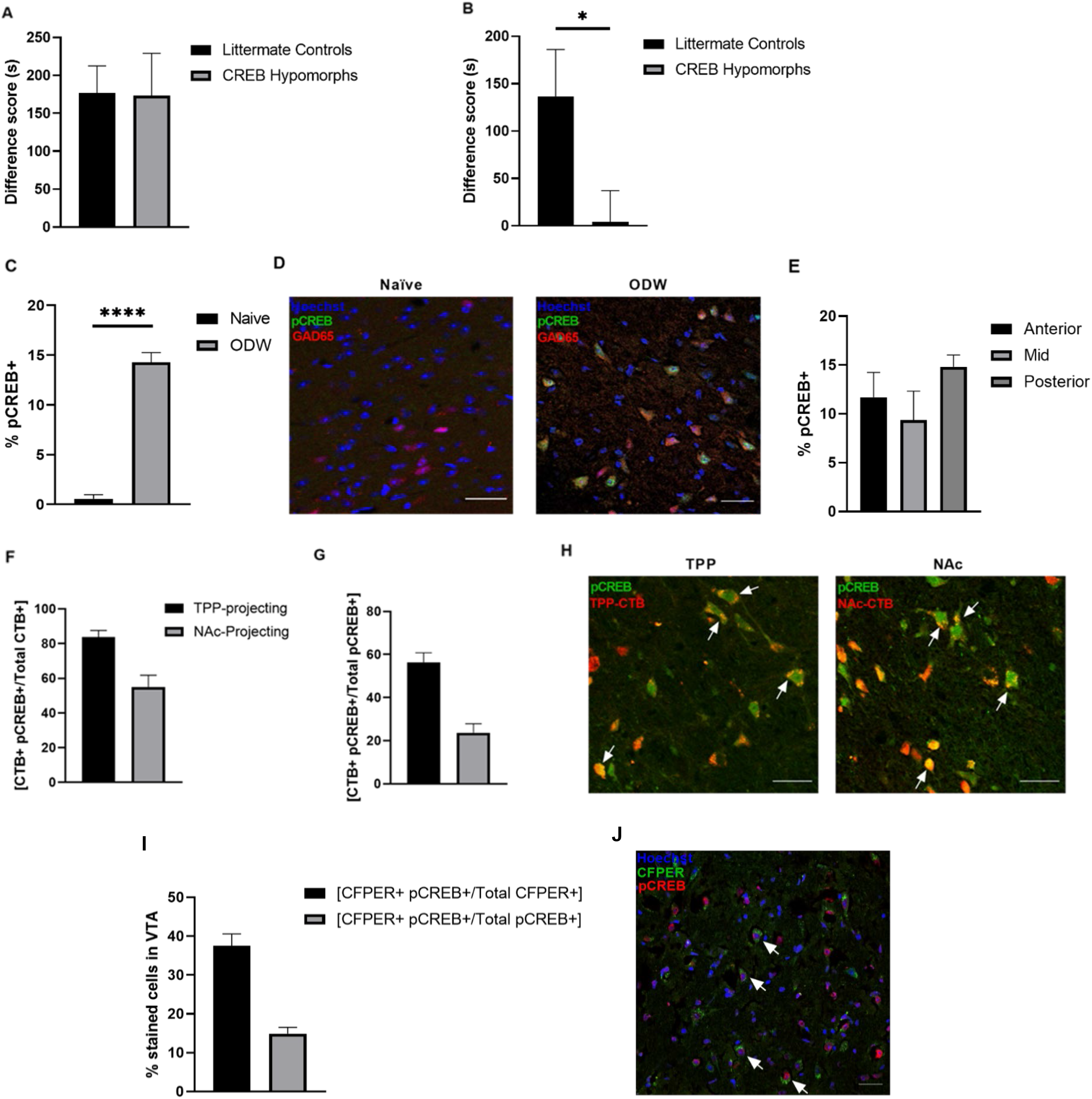
pCREB is upregulated in most TPP-bound VTA neurons of ODW mice. **(A)** Morphine CPP difference scores for drug-naïve CREB homozygous hypomorph mice and littermate controls. Both groups exhibit morphine CPP (two-tailed t-test; *t*_(17)_=0.4763, p=0.9626, *n*=19). **(B)** Morphine CPP difference scores for ODW CREB homozygous hypomorph mice and littermate controls. CREB hypomorphs did not exhibit a morphine CPP, while their littermate counterparts did (two-tailed t-test; *t*(15)=2.162, p=0.0472, *n*=17). Bars represent the mean difference in time spent in the morphine-paired compartment minus the neutral compartment. **(C)** Summary data showing proportion of VTA cells expressing the GABA_A_ switch marker, pCREB in drug-naïve and ODW GAD65-mCherry mice. pCREB expression is significantly higher in ODW mice (*n=*3, 6737 VTA cells) than drug-naïve ones (*n=*3, 2248 VTA cells; Two-tailed *t*_(83)_=11.45, ****: p<0.0001). **(D)** Representative immunofluorescence of pCREB (green) in VTA cells (blue) of drug-naïve and ODW GAD65-mCherry mice (GAD65: red). **(E)** Summary data of anterior, mid, and posterior VTA cells that are pCREB+ in GAD65-mCherry mice. 11.70% of anterior VTA cells, 9.378% of mid VTA cells, and 14.81% of posterior GAD65 stained VTA cells were pCREB+. There was no significant difference in the regional expression of pCREB along the rostrocaudal axis (*F*_(2,40)_=2.280, p=0.115). **(F-H)**Immunohistochemistry of VTA cells in drug-naïve and ODW GAD65-mCherry mice following CTB retrograde tracing from the TPP (*n=*6, 2693 VTA cells) and NAc (*n=*3 mice, 2181 VTA cells). **(F)** Proportion of separate TPP- and NAc-projecting (CTB+) VTA cells that express pCREB. **(G)** Proportion of pCREB+ VTA cells that are TPP-or NAc-projecting. **(H)** Representative immunofluorescence of VTA cells (Hoechst: blue) expressing pCREB (GFP: green) and CTB (red). Arrows: representative double-labeled (yellow) cells with punctate and cytoplasmic staining. Data are represented as mean ± SEM. Scale bars are 30 μm. **(I)** Percentage of stained cells as a fraction of total CFPER+ cells (black bar) and total pCREB+ cells (grey bar). 37% ± 3% of CFPER+ cells co-stained for pCREB, whereas 14% ± 1% of pCREB+ cells co-stained for CFPER. **(J)** Representative immunofluorescence of GAD65-Cre;Cx36fl(CFP)/fl(CFP) VTA cells following staining for CFPER (green), pCREB (red), and Hoechst (blue). Arrows: double-labeled cells. Data are represented as mean ± SEM. ** denotes p<0.01. **** denotes p<0.0001. Scale bars are 30 μm.

Finally, we performed pCREB and CFPER co-staining in a cohort of *GAD65-Cre;Cx36^fl(CFP)/fl(CFP)^* mice that were subjected to the same opiate dependence and withdrawal induction protocol we had used in the GAD65-mCherry mice above. We observed VTA pCREB expression and a large degree of overlap between CFPER+ and pCREB+ neurons wherein almost 40% of CFPER+ neurons were pCREB+ (Figs. 6*I*, *J*). Our observation that a large proportion of Cx36-excised neurons express the functional marker for the GABA_A_ switch, pCREB, raises the possibility that Cx36 contributes to the motivational switch in ODW animals, further warranting functional experiments.

### Pharmacological blockade of VTA Cx36 reverts ODW rats to a drug-naïve motivational state

In order to evaluate the role of VTA Cx36-containing gap junctions in opiate reinforcement, we microinfused the Cx36 blocker, mefloquine, or the vehicle (DMSO) into the VTA of drug-naïve rats and ODW rats prior to morphine CPP training sessions. Intracranial (I.C.) pretreatments were performed 40 minutes prior to conditioning, a timepoint at which mefloquine’s non-specific (i.e., Cx36-independent) effects are largely diminished (Allison et al., 2011). Systemic pretreatments of PBS were administered to all groups in the first iteration of the experiment (Fig. 7*A*). Rats exhibited morphine CPP irrespective of the I.C. pretreatment and motivational state, indicating that VTA mefloquine treatment does not impact the extent to which morphine is reinforcing in drug-naïve or ODW rats (Fig. 7*B*).

**Figure 7.**
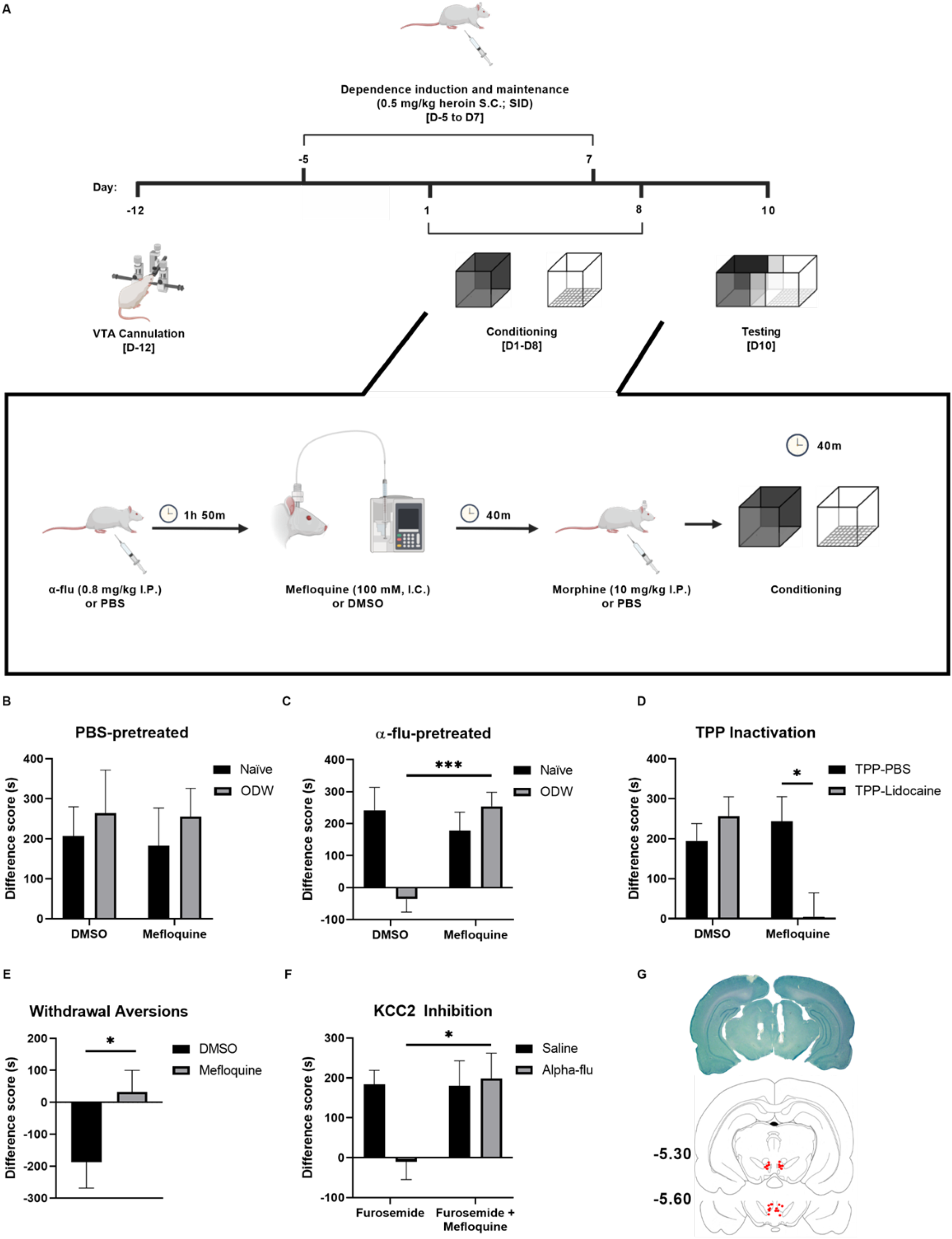
Pharmacological blockade of VTA Cx36 reverts ODW rats to a drug-naïve motivational state. **(A)** Schematic of morphine CPP experiments in rats (created with BioRender.com). **(B-F)** Morphine CPP and opiate withdrawal CPA bar graphs where bars represent the mean difference in time spent in the morphine-paired compartment minus the vehicle-paired compartment, and the mean difference in time spent in the withdrawal-paired compartment minus the neutral compartment, respectively. **(B)** Morphine CPP difference scores for PBS-pretreated drug-naïve and ODW rats receiving intracranial DMSO or mefloquine (*n*=6 per group). CPP difference scores were calculated as time spent in the morphine-paired compartment minus the time spent in the PBS-paired compartment. Rats exhibited morphine CPP irrespective of the I.C. pretreatment and motivational state, as there was no significant effect of motivational state (two-way ANOVA; *F*_(1,20)_=0.545, p=0.468), I.C. pretreatment (*F*_(1,20)_=0.034, p=0.853) nor the motivational state × I.C pretreatment interaction (*F*_(1,20)_=0.008, p=0.929, *n*=24). **(C)** Morphine CPP difference scores for α-flu-pretreated drug-naïve and ODW rats receiving intracranial DMSO or mefloquine (*n*=6 per group). We saw a significant interaction of motivational state × I.C. pretreatment (*F*_(1,20)_=10.26, p=0.0045, *n*=24) where α-flu blocked preferences in DMSO-pretreated ODW rats but not in mefloquine-pretreated ODW rats (two-tailed t-test; *t*_(10)_=4.803, p=0.0007, *n*=12). **(D)** Morphine CPP difference scores for TPP inactivation experiments wherein rats subjected to opiate dependence and withdrawal received PBS or lidocaine into the TPP, and DMSO or mefloquine into the VTA (*n*=30). We saw a significant VTA pretreatment × TPP pretreatment interaction (*F*(1,21)=6.943, p=0.0155, *n*=25). A significant difference was found between rats that received TPP lidocaine and VTA mefloquine vs. rats that received TPP PBS and VTA mefloquine (two-tailed t-test; *t*_(11)_=2.764, p=0.0184, *n*=13). **(E)** CPA difference scores for opiate-withdrawn DMSO and mefloquine-pretreated rats. Unlike the DMSO-pretreated group, mefloquine-pretreated rats did not avoid the withdrawal-paired compartment (two-tailed t-test; *t*_(21)_=2.099, p= 0.0481, *n*=23). **(F)** Morphine CPP difference scores for PBS or α-flu-pretreated rats that either received intracranial furosemide or a cocktail of furosemide and mefloquine. We saw a significant interaction between I.C. pretreatment × systemic pretreatment (*F*_(1,41)_=4.440, p=0.0413) where α-flu blocked CPP in the furosemide group but not in the furosemide + mefloquine group (two-tailed t-test; *t*_(21)_=2.772, p=0.0114, *n*=23). **(G)** Top: representative brain section depicting VTA cannulation following cresyl violet staining. Bottom: schematic of coronal brain sections (−5.30 and −5.60 posterior to bregma). Bilateral cannulae were used and therefore each red dot represents a single cannula tip and has a corresponding red dot on the contralateral side. Subjects were included only if both placements were within the boundaries of the VTA. Schematic depicts placements from mefloquine-pretreated ODW rats in Fig 3c. Drawings adapted from Paxinos & Watson (Paxinos and Watson, 2006). Data are represented as mean ± SEM. * denotes differences where p<0.05. ** denotes differences where p<0.01.

To assess whether VTA Cx36 is implicated in the switch to DA-mediated ODW motivation, we performed the same experiment but with systemic α-flupenthixol (α-flu) pretreatments in separate cohorts of naïve rats and ODW rats. Consistent with evidence indicating that drug-naïve reinforcement is DA-independent (Bechara and van der Kooy, 1992), α-flu had no effect on morphine CPP in both drug-naïve groups (i.e. DMSO and mefloquine-pretreated). As expected, α-flu blocked morphine CPP in DMSO-pretreated ODW rats. By contrast, α-flu did not abolish morphine CPP in mefloquine-pretreated ODW rats (Fig. 7*C*), suggesting that Cx36 blockade in the VTA prevents the manifestation of a DA-mediated ODW motivational state. We hypothesized that this may occur through a reversion of ODW rats to a TPP-mediated, drug-naïve motivational state. To test this, we reversibly inactivated the TPP with lidocaine in mefloquine-pretreated rats subjected to the same opiate dependence and withdrawal induction protocol. TPP lidocaine blocked morphine CPP in ODW rats that received VTA mefloquine infusions (Fig. 7*D*). These results support the notion that blockade of VTA Cx36 in ODW rats causes a reversion to a TPP-mediated, drug-naïve motivational state.

If mefloquine-pretreated ODW rats are being reverted to a drug-naïve motivational state, they should not exhibit opiate withdrawal aversions following chronic opiate cessation. We performed a CPA paradigm and found that, unlike the DMSO-pretreated control group, mefloquine-pretreated ODW rats did not avoid the withdrawal-paired compartment (Fig. 7*E*).

Previous work has demonstrated that inhibiting the K-Cl cotransporter, KCC2, in the VTA results in a switch from a drug-naïve motivational state to an ODW one – the inverse effect of Cx36 blockade (Ting-A-Kee et al., 2013). In other words, VTA infusions of the KCC2 inhibitor, furosemide, in drug-naïve rats result in a behavioural and electrophysiological phenocopying of the ODW motivational state in opiate-naïve animals. This raises the possibility that Cx36 and KCC2 are acting through the same signaling pathway. To this end, we microinfused a cocktail of mefloquine and furosemide to see which effect would persist over the other, an indirect measure of whether Cx36 is functioning up- or downstream of KCC2. Neither furosemide nor the cocktail of furosemide and mefloquine interfered with the ability of PBS-pretreated drug-naïve rats to exhibit morphine CPP. Consistent with previous reports (Ting-A-Kee et al., 2013), morphine CPP was abolished by systemic α-flu in the furosemide-pretreated ODW group. By contrast, α-flu did not abolish morphine CPP in ODW rats pretreated with the mefloquine and furosemide cocktail (Fig. 7*F*). The dominance of the mefloquine effect over furosemide suggests that Cx36 functions downstream of KCC2.

Our observations that VTA mefloquine reverts ODW rats to a drug-naïve state wherein morphine CPP is DA-independent, TPP-dependent, and withdrawal aversions are abolished, suggest that VTA Cx36 is necessary for the development of a DA-mediated ODW motivational state. Moreover, the effect of intra-VTA mefloquine cannot be explained by a generalized block of aversions as systemic lithium chloride elicited CPA in both DMSO and mefloquine-pretreated rats (Fig. 8*A*). Mefloquine did not impact somatic withdrawal signs, indicating a specific effect on affective withdrawal (Fig. 8*B*).

**Figure 8.**
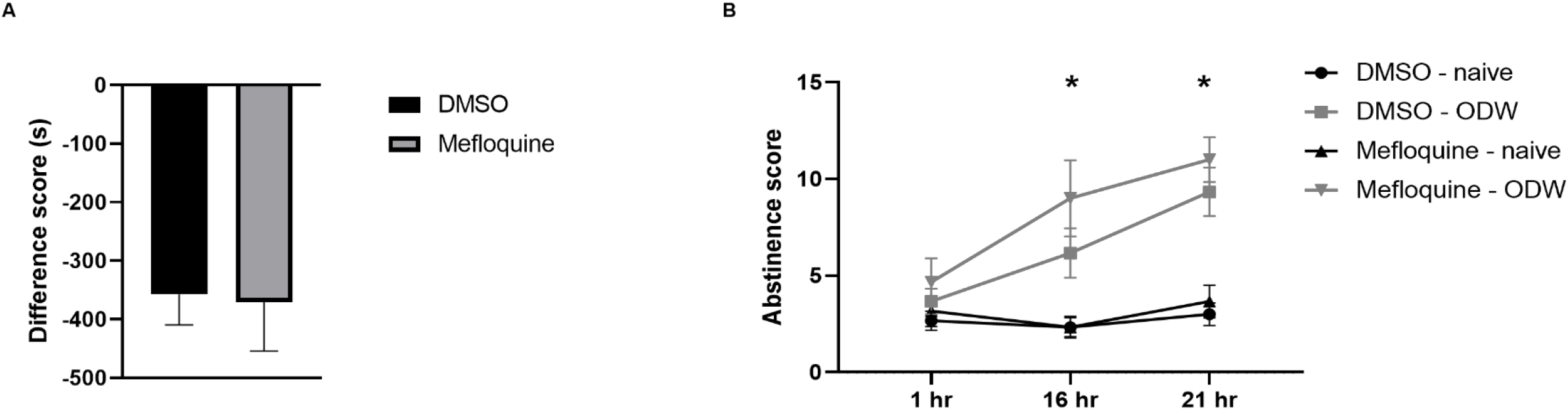
Pharmacological blockade of VTA Cx36 spares LiCl CPA and somatic opiate withdrawal signs in rats. **(A)** Rats spent less time in a previously LiCl-paired compartment compared to a previously vehicle-paired compartment irrespective of whether they were DMSO-or mefloquine-pretreated (two-tailed t-test; t(6)=0.1387, p=0.8942; *n*=8). Bars represent the mean difference in time spent in the LiCl-paired compartment minus the vehicle-paired compartment. Error bars represent ± SEM. **(B)** Drug-naïve mice exhibited no somatic withdrawal signs at any timepoint following the last heroin injection, which was administered after a 5-day period of heroin administration. By contrast, ODW mice did exhibit somatic signs16 and 21 hours following the last heroin administration, irrespective of whether they were DMSO-or mefloquine-pretreated. As such, we saw significant effects of time (*F*_(2,60)_=9.669, p=0.0002), I.C. pretreatment (*F*_(3,60)_=20.11, p<0.0001), and time × pretreatment (*F*_(6,60)_=2.734, p=0.0205; *n*=24). Bonferroni’s tests revealed a significant difference at 16 hrs and 21 hrs between DMSO-pretreated drug-naïve rats and DMSO-pretreated ODW rats (*n*=12, p=0.0003), DMSO-pretreated drug-naïve rats and mefloquine-pretreated ODW rats (*n*=12, p<0.0001), mefloquine-pretreated drug-naïve rats and mefloquine-pretreated ODW rats (*n*=12, p<0.0001), and mefloquine-pretreated drug-naïve rats and DMSO-pretreated ODW rats (*n*=12, p=0.0013). There was no difference between the two ODW groups. There also was no difference between the drug-naïve groups.

### GAD65-Cre;Cx36^*fl(CFP)/fl(CFP)*^ mice are perpetually drug-naïve

To rule out non-specific effects of mefloquine, we performed a similar set of morphine CPP (Fig. 9*A*) and withdrawal CPA (Fig. 9*D*) experiments using *GAD65-Cre;Cx36^fl(CFP)/fl(CFP)^* mice. When pretreated with systemic PBS, *GAD65-Cre;Cx36^fl(CFP)/fl(CFP)^* mice exhibited morphine CPP to the same extent as littermate controls irrespective of whether they were drug-naïve or ODW (Fig. 9*B*). By contrast, systemic α-flu abolished ODW morphine CPP in littermate controls but not in *GAD65-Cre;Cx36^fl(CFP)/fl(CFP)^* mice (Fig. 9*C*). The inability of α-flu to block morphine CPP in *GAD65-Cre;Cx36^fl(CFP)/fl(CFP)^* mice subjected to chronic opiates suggests that they are in a drug-naïve motivational state mediated by the TPP. If so, they should not exhibit a CPA in response to spontaneous opiate withdrawal following opiate dependence induction. Indeed, although ODW littermate controls avoided the withdrawal-paired compartment, *GAD65-Cre;Cx36^fl(CFP)/fl(CFP)^*mice did not (Fig. 9*E*). *GAD65-Cre;Cx36^fl(CFP)/fl(CFP)^* mice were more likely to exhibit this drug-naïve-like behaviour – an indifference to the two compartments – if they had a higher number of VTA neurons expressing the excision reporter in the VTA (Figs. 9*F*, *G*). The lack of withdrawal aversions cannot be explained by a generalized block of aversions, as LiCl elicited CPA in both groups of mice (Fig. 10).

**Figure 9.**
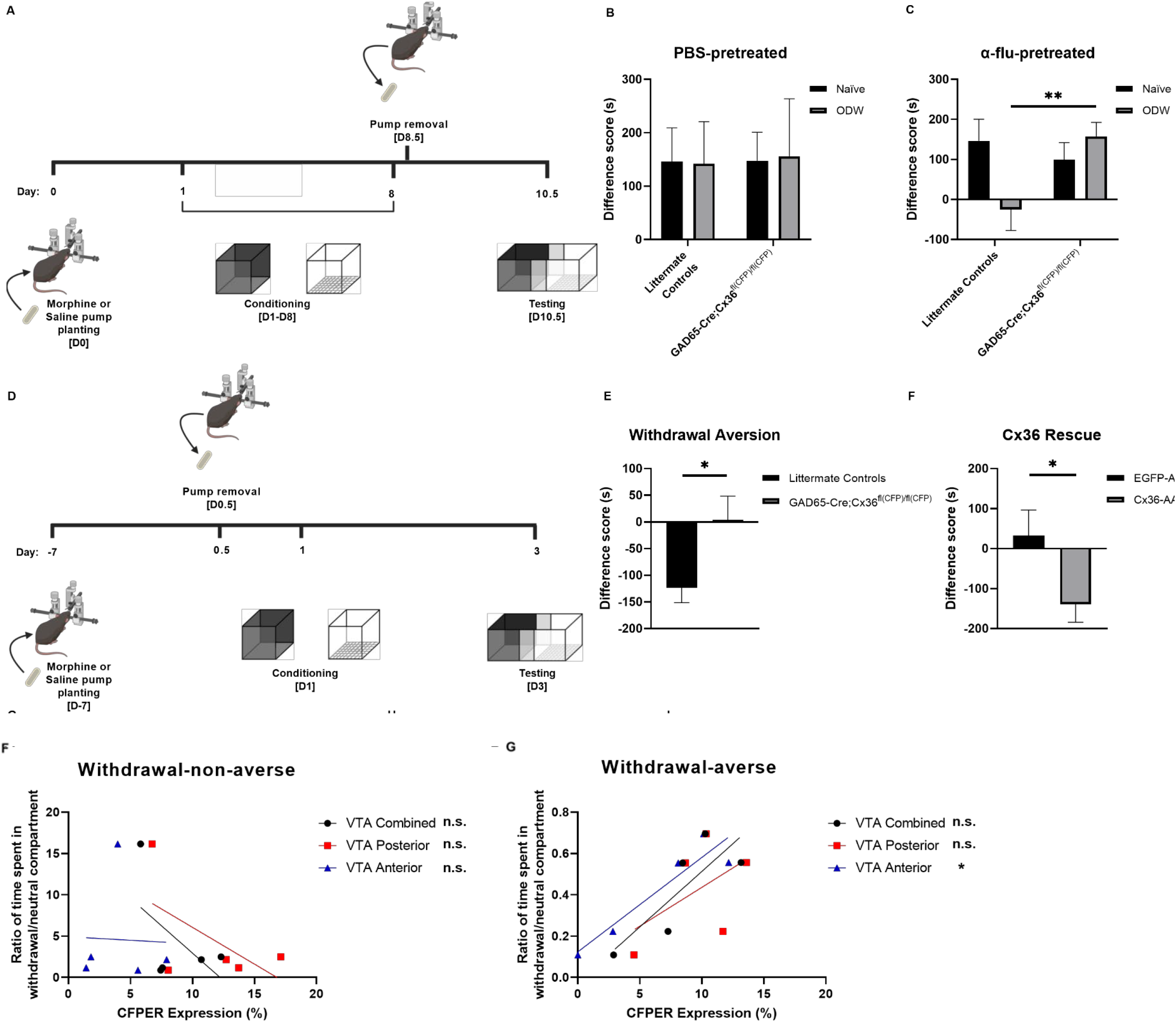
*GAD65-Cre;Cx36^fl(CFP)/fl(CFP)^* mice are perpetually drug-naïve. **(A)** Schematic of morphine CPP experiments in mice (created with BioRender.com). **(B-C)** Morphine CPP bar graphs where bars represent the mean difference in time spent in the morphine-paired compartment minus the vehicle-paired compartment. **(B)** Morphine CPP difference scores for saline-pretreated drug-naïve and ODW *GAD65-Cre;Cx36^fl(CFP)/fl(CFP)^* mice and littermate controls *GAD65-Cre;Cx36^fl(CFP)/fl(CFP^* mice exhibited morphine CPP to the same extent as littermate controls irrespective of whether they were drug-naïve or ODW as we saw no main effect of genotype (*F*_(1,32)_=0.0007656, p=0.9781), motivational state (*F*_(1, 32)_=0.008424, p=0.9274) nor an interaction (*F*_(1,32)_=0.005017, p=0.9440; *n*=36). **(C)** Morphine CPP difference scores for α-flu-pretreated drug-naïve and ODW *GAD65-Cre;Cx36^fl(CFP)/fl(CFP^* mice and littermate controls. *GAD65-Cre;Cx36^fl(CFP)/fl(CFP^* mice exhibited morphine CPP irrespective of whether they were drug-naïve or ODW. We saw a significant genotype × motivational state interaction (*F*_(1,53)_=5.776, p=0.0198; *n*=57) where α-flu blocked preferences in ODW littermate controls but not ODW *GAD65-Cre;Cx36^fl(CFP)/fl(CFP)^* mice (one-tailed t-test; t_(25)_=2.780, p=0.0051, *n*=27). **(D)** Schematic for morphine withdrawal CPA experiments in mice (created with BioRender.com). **(E)** CPA difference scores for opiate-withdrawn *GAD65-Cre;Cx36^fl(CFP)/fl(CFP^* mice and littermate controls. Although littermate controls avoided the withdrawal-paired compartment, *GAD65-Cre;Cx36^fl(CFP)/fl(CFP^* mice did not (one-tailed t-test; *t*_(28)_=2.401, p=0.0116; *n*=30). **(F)** Chronic morphine withdrawal CPA data plotted against VTA CFPER expression (i.e. % CFPER+ in the VTA) data from the five *GAD65-Cre;Cx36^fl(CFP)/fl(CFP)^* mice that spent the *most* time in the withdrawal-paired compartment (withdrawal-non-averse). Among those five mice, CFPER expression was highest in those that liked withdrawal the least and therefore exhibited the most drug-naïve-like phenotype (indifference to the two compartments), akin to the majority of *GAD65-Cre;Cx36^fl(CFP)/fl(CFP)^*. This trend was not significant: R^2^_(VTA Combined)_ = 0.2910, p_(VTA Combined)_ = 0.3481; R^2^_(VTA Posterior)_ = 0.3389, p_(VTA Posterior)_ = 0.3031; and R^2^_(VTA Anterior)_ = 0.001232, p_(VTA Anterior)_ = 0.9553. **(G)** Chronic morphine withdrawal CPA data plotted against VTA CFPER expression data from the five *GAD65-Cre;Cx36^fl(CFP)/fl(CFP)^* mice that spent the *least* time in the withdrawal-paired compartment (withdrawal-averse). Among those five mice, CFPER expression was highest in those that avoided withdrawal the least and thus exhibited the most drug-naïve-like phenotype (indifference to the two compartments). This correlation was significant in the anterior VTA but not in the posterior and entirety of the VTA: R^2^_(VTA Combined)_ = 0.6745, p_(VTA Combined)_ = 0.0882; R^2^_(VTA Posterior)_ = 0.2707, p_(VTA Posterior)_ = 0.3688; R^2^_(VTA Anterior)_ = 0.8724, p_(VTA Anterior)_ = 0.0201. * denotes p<0.05.

**Figure 10.**
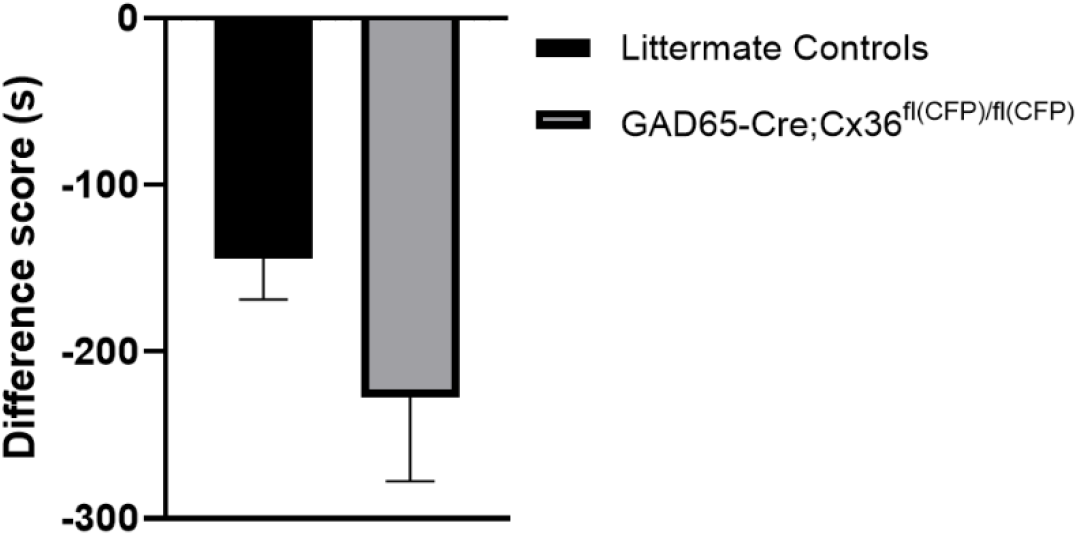
Genetic ablation of Cx36 in GAD65+ cells spares LiCl CPA. No difference in LiCl CPA between *GAD65-Cre;Cx36^fl(CFP)/fl(CFP)^* mice and littermate controls, as both groups spent less time in the LiCl-paired compartment compared to a vehicle-paired compartment (two-tailed t-test; t_(14)_=1.476, p=0.1621; *n*=16). Bars represent the mean difference in time spent in the LiCl-paired compartment minus the vehicle-paired compartment. Error bars represent ± SEM.

These experiments corroborate the findings from the pharmacological work in rats – that Cx36 is necessary for the manifestation of ODW motivation. However, neither our pharmacological manipulations in rats nor our genetic manipulations in mice achieve regional and cell-type specificity simultaneously. Therefore, we performed a Cx36 rescue experiment in which we injected a Cre-dependent adeno-associated virus expressing Cx36 (Cx36-AAV) or a control virus (EGFP-AAV) into the VTA of *GAD65-Cre;Cx36^fl(CFP)/fl(CFP)^* mice. In so doing, Cx36 expression is rescued exclusively in the VTA GABA neurons of the Cx36-AAV group. The Cx36-AAV group exhibited CPA in response to opiate withdrawal whereas the EGFP-AAV group did not (Fig. 11*A*). Immunostaining revealed that almost 20% of VTA cells were Cx36+ in the Cx36-AAV group, whereas only about 7% ± 1% were Cx36+ in the EGFP-AAV group (Fig. 11*B*, *C*). This is perhaps due to Cx36 expression in GAD67+ GABA neurons that lack GAD65 or inefficient Cre-lox recombination (Taylor et al., 2014). This illustrates that Cx36 rescue in VTA GABA neurons is sufficient to restore the susceptibility of *GAD65-Cre;Cx36^fl(CFP)/fl(CFP)^* mice to ODW motivation.

**Figure 11.**
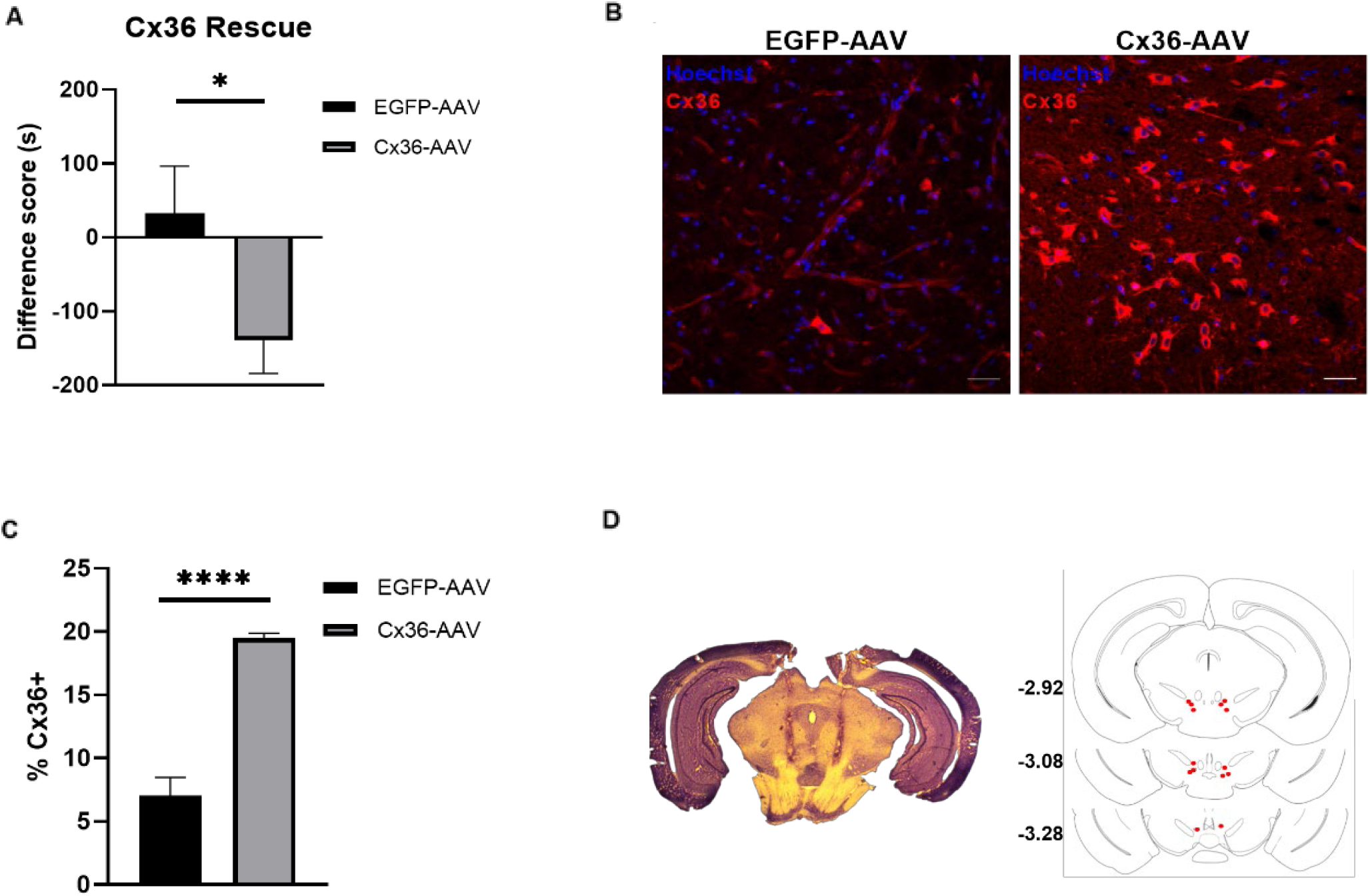
Rescue of Cx36 in VTA GABA neurons of *GAD65-Cre;Cx36fl(CFP)/fl(CFP)* mice restores their susceptibility to opiate withdrawal aversions. **(A)** CPA difference scores for opiate-withdrawn *GAD65-Cre;Cx36^fl(CFP)/fl(CFP^* mice that received intra-VTA injections of a Cre-dependent adeno-associated virus expressing Cx36 (Cx36-AAV) or a control virus (EGFP-AAV). The Cx36-AAV group exhibited CPA in response to opiate-withdrawal whereas the EGFP-AAV group did not (one-tailed t-test; *t*_(25)_=2.159, p=0.0203; *n*=27). **(B)** Representative immunofluorescence of *GAD65-Cre;Cx36^fl(CFP)/fl(CFP^* VTA transfected with EGFP-AAV (left) or Cx36-AAV (right) and stained using a Cx36 antibody (red). Nuclei were stained with Hoechst (blue). **(C)** Summary data of VTA neurons expressing Cx36 in *GAD65-Cre;Cx36^fl(CFP)/fl(CFP^* mice that received EGFP-AAV or Cx36-AAV in the VTA. Cx36 expression is significantly higher in mice that received the Cx36 rescue construct (19.52%; *n*=4 mice, 10381 VTA cells) compared to those that received the control construct (7.053%; *n*=4 mice, 4789 VTA cells; one-tailed t-test; *t*_(112)_=11.35, ****: p<0.0001). **(D)** Left: Representative mouse brain section stained with cresyl violet to visualize injection track targeting VTA for viral transfection. Right: schematic of coronal brain sections (−2.92, −3.08 and −3.28 posterior to bregma). Viral injections were bilateral and therefore each red dot represents a single injector tip and has a corresponding red dot representing the contralateral injection in the same subject. If either injector missed the VTA, the subject was excluded. Schematic depicts viral injection placements from Cx36-AAV group in Fig 4f. Drawings adapted from Allen Mouse Brain Atlas (Lein et al., 2007). Data are represented as mean ± SEM. * denotes differences where p<0.05. ** denotes differences where p<0.01. **** denotes differences where p<0.0001. Scale bars are 30 μm.

## Discussion

We show that VTA Cx36 is upregulated upon opiate dependence and withdrawal, that pharmacological blockade of VTA Cx36 reverts ODW rats to a drug-naïve motivational state, that mice lacking Cx36 in GABAergic neurons are perpetually drug-naïve with respect to motivation, and that the susceptibility of those mice to opiate dependence is restored when Cx36 is rescued exclusively in VTA GABA neurons. Put together, this work suggests that Cx36 in VTA GABA neurons is both necessary for the manifestation of opiate-dependent and withdrawn motivation and sufficient for imparting susceptibility to that very dependence. Chronic opiates and withdrawal cause an upregulation of BDNF in the VTA, which has been shown to downregulate KCC2 expression thereby triggering a positive shift in the GABA_A_ reversal potential, thus producing excitatory effects of GABA_A_ agonists (Rivera et al., 2002; Vargas-Perez et al., 2009). GABA_A_ signaling has robust modulatory effects on Cx36 expression and electrical coupling early in development and thus it is conceivable that a similar mechanism is at play in the VTA under ODW conditions, although a direct interplay remains to be demonstrated (Park et al., 2011). Our finding that the ‘naïve-state-inducing’ effect of Cx36 blockade dominates over the ‘dependence-inducing’ effect of KCC2 inhibition suggests that Cx36 acts downstream of KCC2 with respect to the opiate motivational switch. In addition, although *GAD65-Cre;Cx36^fl(CFP)/fl(CFP)^* mice never transition to an ODW motivational state, they still exhibit the VTA pCREB staining that is characteristic of ODW VTA. This suggests that pCREB may be acting as an upstream effector of Cx36 – an idea not without precedent as there is evidence of CREB-mediated regulation of Cx36 gap junctional uncoupling early in development (Arumugam et al., 2005).

In ODW animals, pCREB is upregulated in VTA GABA neurons with excitatory GABA_A_ receptors (Laviolette et al., 2004). Our finding that the vast majority of TPP-projecting cells are pCREB+ suggests that the TPP receives predominant input from hyperexcitable (i.e. ‘switched’) VTA GABA neurons. We hypothesize that this projection masks the TPP-mediated, drug-naïve motivational state in ODW animals. Two lines of evidence inform this hypothesis: 1) re-expressing presynaptic μ-opiate receptor (μOR) on NAc terminals in the VTA is sufficient to restore opiate motivation in otherwise μOR KO mice, indicating that the motivational effects of opiates are likely produced via presynaptic mechanisms (Cui et al., 2014). VTA GABA neurons with excitatory GABA_A_ receptors would be inhibited by presynaptic opiate action. 2) GABA release in the TPP is a prerequisite for morphine reinforcement in drug-naïve animals as blocking GABAB receptors in the TPP blocks morphine CPP in drug-naïve rats (Heinmiller et al., 2009). Our speculation is that the opiate-induced inhibition of hyperexcitable, TPP-projecting GABA neurons shuts down the TPP, thereby rendering it functionally dispensable during opiate withdrawal (Fig. 12).

**Figure 12.**
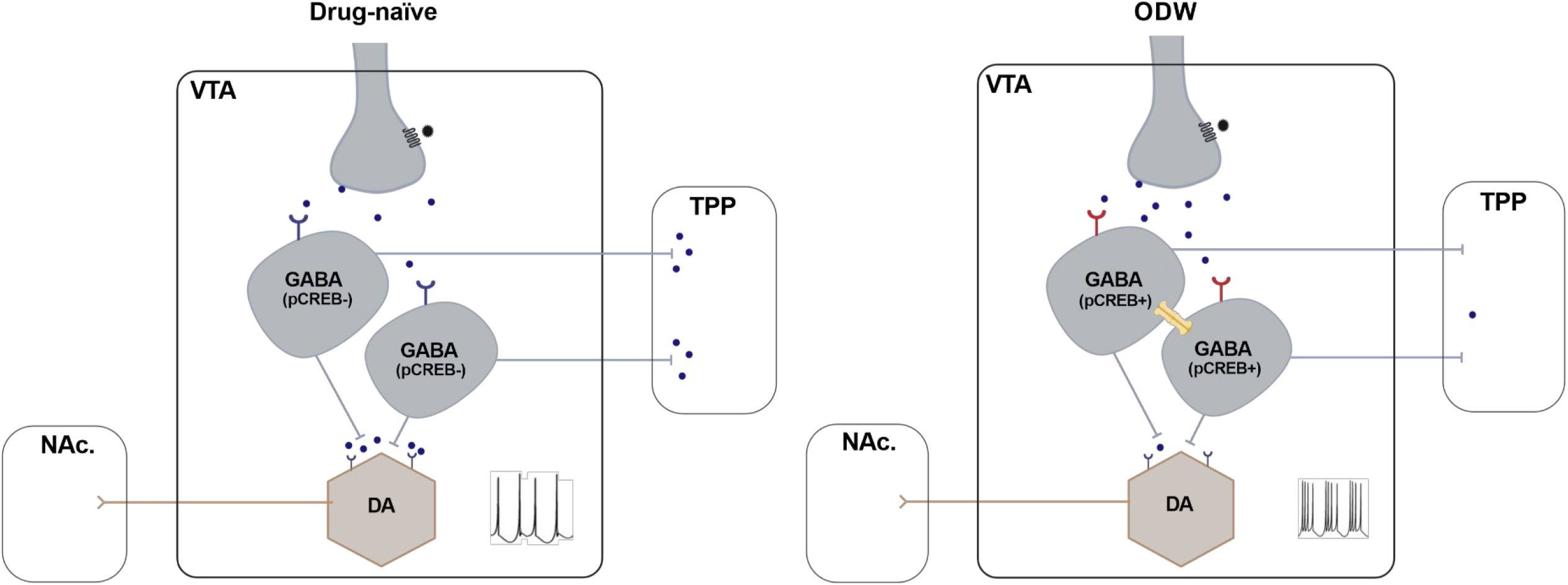
Proposed mechanisms of opiate reinforcement in drug-naïve and ODW VTA. Schematics depicting the VTA and its outputs following morphine administration under two conditions: a drug-naïve (left) and ODW (right) motivational state. Left: The binding of morphine onto presynaptic mu opiate receptors results in decreased GABA release onto VTA GABA neurons. In drug-naïve conditions, VTA GABA neurons are ‘unswitched’ (i.e., express inhibitory GABAA receptors and do not express pCREB) and, as such, GABA-GABAA binding elicits a hyperpolarizing response. Therefore, the morphine-induced decrease in GABA release likely results in VTA GABA neuron disinhibition and a consequent increase in GABA release from descending projections to the TPP and from local collaterals onto VTA DA neurons. GABA release in the TPP is necessary for TPP-mediated mCPP, therefore, we speculate that this projection is functionally implicated in opiate reinforcement exhibited by drug-naïve animals. In addition, the increased release of GABA onto VTA DA neurons may serve as an ‘OFF’ switch for the mesolimbic DA system that is necessary for ODW opiate reinforcement. Right: In ODW VTA, the binding of opiates onto presynaptic mu opiate receptors also results in decreased GABA release onto VTA GABA neurons. However, in this case, given that half of the VTA GABA population is ‘switched’ (i.e., expresses excitatory GABAA receptors and pCREB), the morphine-induced decrease in GABA release would inhibit any switched VTA GABA neurons. The vast majority of TPP-projecting GABA neurons from the VTA are pCREB+ in ODW animals and, therefore, we speculate that there is a reduction of GABA release in the TPP following opiate administration in ODW animals. Drug-naïve mCPP requires GABA-GABAB binding in the TPP so we predict that a reduction of GABA release from the VTA to the TPP may serve as an ‘OFF’ switch for the TPP system under ODW conditions. In addition, Cx36 gap VTA GABA neuron junctions may mediate an ‘ON’ switch for the mesolimbic DA system, as other lines of work have shown that increased synchrony and coupling of VTA GABA neurons results in increased DA burst firing (Morozova et al., 2016).

Our behavioural experiments did not dissect the functional contributions of Cx36 in a projection-specific manner. Therefore, it remains unclear which subpopulation of Cx36-expressing neurons is functionally important in the switch to a DA-dependent, ODW motivational state. *In vivo* extracellular recordings in anesthetized rats reveal an increase in DA burst firing in response to increased VTA GABA synchrony (Morozova et al., 2016). Such synchronous activity is a hallmark feature of neuronal networks coupled by electrical synapses composed of Cx36-containing gap junctions (Nagy et al., 2018; Pereda, 2014; Pereda et al., 2013). We therefore speculate that interneuronal Cx36 contributes to the motivational switch via DA neuron disinhibition and a consequent activation of the DA-mediated, ODW motivational state.

Our findings identify VTA gap junctions as key regulators of drug reinforcement that have yet to be incorporated into prevailing circuit models of addiction. The motivational switch from reward-seeking to withdrawal-avoidance is a hallmark of substance use disorders and appears to be determined by Cx36. Understanding the mechanisms by which Cx36 dictates motivation and produces withdrawal avoidance is a pivotal step toward novel treatments. Moreover, the preponderance of quinolone drugs that inhibit gap junctions begs further investigation of their therapeutic potential.

## Acknowledgements

This work was supported by the Canadian Institutes of Health Research (CIHR). We would like to thank the Division of Comparative Medicine at the University of Toronto for animal colony husbandry and maintenance, and Dr. Kate Banks and Dr. Chereen Collymore for their continual guidance on surgeries and animal welfare.

## Author contributions

G.M., under the supervision and guidance of D.v.d.K., wrote the manuscript and designed, performed, and analyzed the majority of surgical procedures and behavioural experiments. M.Y., M.B., M.G., R.C., E.K., S.D.V., and M.A. performed stereotaxic surgeries and performed some behavioural and immunohistological work and associated analyses. E.H., under the supervision of R.B., designed, conducted, and analyzed experiments characterizing mouse strain used for tracing. T.G. provided expertise in design of mouse CPP paradigm. J.N. supplied Cx36-EGFP mouse strain, provided protocols and samples for histological work and shared his expertise on Cx36.

The authors declare no competing interests.

## References

1. Allison, D.W., Ohran, A.J., Stobbs, S.H., Mameli, M., Valenzuela, C.F., Sudweeks, S.N., Ray, A.P., Henriksen, S.J., and Steffensen, S.C. (2006). Connexin-36 gap junctions mediate electrical coupling between ventral tegmental area GABA neurons. Synapse 60, 20–31.

2. Allison, D.W., Wilcox, R.S., Ellefsen, K.L., Askew, C.E., Hansen, D.M., Wilcox, J.D., Sandoval, S.S., Eggett, D.L., Yanagawa, Y., and Steffensen, S.C. (2011). Mefloquine effects on ventral tegmental area dopamine and GABA neuron inhibition: A physiologic role for connexin-36 gap junctions. Synapse 65, 804–813.

3. Arumugam, H., Liu, X., Colombo, P.J., Corriveau, R.A., and Belousov, A.B. (2005). NMDA receptors regulate developmental gap junction uncoupling via CREB signaling. Nature Neuroscience 8, 1720–1726.

4. Barmashenko, G., Hefft, S., Aertsen, A., Kirschstein, T., and Köhling, R. (2011). Positive shifts of the GABA_A_ receptor reversal potential due to altered chloride homeostasis is widespread after status epilepticus: Positive Shifts of the GABA_A_ Receptor Reversal Potential. Epilepsia 52, 1570–1578.

5. Bechara, A., and van der Kooy, D. (1992). A single brain stem substrate mediates the motivational effects of both opiates and food in nondeprived rats but not in deprived rats. Behavioral Neuroscience 106, 351–363.

6. Bissiere, S., Zelikowsky, M., Ponnusamy, R., Jacobs, N.S., Blair, H.T., and Fanselow, M.S. (2011). Electrical Synapses Control Hippocampal Contributions to Fear Learning and Memory. Science 331, 87–91.

7. Coull, J.A.M., Beggs, S., Boudreau, D., Boivin, D., Tsuda, M., Inoue, K., Gravel, C., Salter, M.W., and De Koninck, Y. (2005). BDNF from microglia causes the shift in neuronal anion gradient underlying neuropathic pain. Nature 438, 1017–1021.

8. Cui, Y., Ostlund, S.B., James, A.S., Park, C.S., Ge, W., Roberts, K.W., Mittal, N., Murphy, N.P., Cepeda, C., Kieffer, B.L., et al. (2014). Targeted expression of μ-opioid receptors in a subset of striatal direct-pathway neurons restores opiate reward. Nature Neuroscience 17, 254–261.

9. Deans, M.R., Gibson, J.R., Sellitto, C., Connors, B.W., and Paul, D.L. (2001). Synchronous activity of inhibitory networks in neocortex requires electrical synapses containing connexin36. Neuron 31, 477–485.

10. Dockstader, C.L., Rubinstein, M., Grandy, D.K., Low, M.J., and Kooy, D.V.D. (2001). The D2 receptor is critical in mediating opiate motivation only in opiate-dependent and withdrawn mice. European Journal of Neuroscience 13, 995–1001.

11. Franco-Pérez, J., Ballesteros-Zebadúa, P., and Manjarrez-Marmolejo, J. (2015). Anticonvulsant effects of mefloquine on generalized tonic-clonic seizures induced by two acute models in rats. BMC Neurosci 16, 7.

12. Galarreta, M., and Hestrin, S. (2001). Electrical synapses between GABA-releasing interneurons. Nat. Rev. Neurosci. 2, 425–433.

13. Heinmiller, A., Ting-A-Kee, R., Vargas-Perez, H., Yeh, A., and van der Kooy, D. (2009). Tegmental pedunculopontine glutamate and GABA-B synapses mediate morphine reward. Behavioral Neuroscience 123, 145–155.

14. Koob, G.F., and Volkow, N.D. (2010). Neurocircuitry of Addiction. Neuropsychopharmacology 35, 217–238.

15. Laviolette, S.R., and van der Kooy, D. (2001). GABA A receptors in the ventral tegmental area control bidirectional reward signalling between dopaminergic and non-dopaminergic neural motivational systems: GABAergic reward signalling in the ventral tegmental area. European Journal of Neuroscience 13, 1009–1015.

16. Laviolette, S.R., Alexson, T.O., and van der Kooy, D. (2002). Lesions of the Tegmental Pedunculopontine Nucleus Block the Rewarding Effects and Reveal the Aversive Effects of Nicotine in the Ventral Tegmental Area. The Journal of Neuroscience 22, 8653–8660.

17. Laviolette, S.R., Gallegos, R.A., Henriksen, S.J., and van der Kooy, D. (2004). Opiate state controls bi-directional reward signaling via GABA_A_ receptors in the ventral tegmental area. Nature Neuroscience 7, 160–169.

18. Lein, E.S., Hawrylycz, M.J., Ao, N., Ayres, M., Bensinger, A., Bernard, A., Boe, A.F., Boguski, M.S., Brockway, K.S., Byrnes, E.J., et al. (2007). Genome-wide atlas of gene expression in the adult mouse brain. Nature 445, 168–176.

19. Margolis, E.B., Toy, B., Himmels, P., Morales, M., and Fields, H.L. (2012). Identification of Rat Ventral Tegmental Area GABAergic Neurons. PLoS ONE 7, e42365.

20. Morozova, E.O., Myroshnychenko, M., Zakharov, D., di Volo, M., Gutkin, B., Lapish, C.C., and Kuznetsov, A. (2016). Contribution of synchronized GABAergic neurons to dopaminergic neuron firing and bursting. J Neurophysiol 116, 1900–1923.

21. Mucha, R.F., van der Kooy, D., O’Shaughnessy, M., and Bucenieks, P. (1982). Drug reinforcement studied by the use of place conditioning in rat. Brain Research 243, 91–105.

22. Nader, K., and van der Kooy, D. (1997). Deprivation State Switches the Neurobiological Substrates Mediating Opiate Reward in the Ventral Tegmental Area. The Journal of Neuroscience 17, 383–390.

23. Nader, K., Bechara, A., Roberts, D.C.S., and van der Kooy, D. (1994). Neuroleptics block high- but not low-dose heroin place preferences: Further evidence for a two-system model of motivation. Behavioral Neuroscience 108, 1128–1138.

24. Nagy, J.I., Pereda, A.E., and Rash, J.E. (2018). Electrical synapses in mammalian CNS: Past eras, present focus and future directions. Biochimica et Biophysica Acta (BBA)-Biomembranes 1860, 102–123.

25. Nair-Roberts, R.G., Chatelain-Badie, S.D., Benson, E., White-Cooper, H., Bolam, J.P., and Ungless, M.A. (2008). Stereological estimates of dopaminergic, GABAergic and glutamatergic neurons in the ventral tegmental area, substantia nigra and retrorubral field in the rat. Neuroscience 152, 1024–1031.

26. Olmstead, M.C., Munn, E.M., Franklin, K.B.J., and Wise, R.A. (1998). Effects of Pedunculopontine Tegmental Nucleus Lesions on Responding for Intravenous Heroin under Different Schedules of Reinforcement. The Journal of Neuroscience 18, 5035–5044.

27. Park, W.-M., Wang, Y., Park, S., Denisova, J.V., Fontes, J.D., and Belousov, A.B. (2011). Interplay of Chemical Neurotransmitters Regulates Developmental Increase in Electrical Synapses. J Neurosci 31, 5909–5920.

28. Paxinos, G., and Watson, C. (2006). The Rat Brain in Stereotaxic Coordinates: Hard Cover Edition (Elsevier).

29. Pereda, A.E. (2014). Electrical synapses and their functional interactions with chemical synapses. Nat Rev Neurosci 15, 250–263.

30. Pereda, A.E., Curti, S., Hoge, G., Cachope, R., Flores, C.E., and Rash, J.E. (2013). Gap junction-mediated electrical transmission: Regulatory mechanisms and plasticity. Biochimica et Biophysica Acta (BBA) - Biomembranes 1828, 134–146.

31. Peron, S.P., Freeman, J., Iyer, V., Guo, C., and Svoboda, K. (2015). A Cellular Resolution Map of Barrel Cortex Activity during Tactile Behavior. Neuron 86, 783–799.

32. Rivera, C., Li, H., Thomas-Crusells, J., Lahtinen, H., Viitanen, T., Nanobashvili, A., Kokaia, Z., Airaksinen, M.S., Voipio, J., Kaila, K., et al. (2002). BDNF-induced TrkB activation down-regulates the K+–Cl− cotransporter KCC2 and impairs neuronal Cl− extrusion. J Cell Biol 159, 747–752.

33. Swanson, L.W. (1982). The projections of the ventral tegmental area and adjacent regions: A combined fluorescent retrograde tracer and immunofluorescence study in the rat. Brain Research Bulletin 9, 321–353.

34. Taniguchi, H., He, M., Wu, P., Kim, S., Paik, R., Sugino, K., Kvitsani, D., Fu, Y., Lu, J., Lin, Y., et al. (2011). A Resource of Cre Driver Lines for Genetic Targeting of GABAergic Neurons in Cerebral Cortex. Neuron 71, 995–1013.

35. Taylor, S.R., Badurek, S., Dileone, R.J., Nashmi, R., Minichiello, L., and Picciotto, M.R. (2014). GABAergic and glutamatergic efferents of the mouse ventral tegmental area: Mouse VTA projections. J. Comp. Neurol. 522, 3308–3334.

36. Ting-A-Kee, R., Dockstader, C., Heinmiller, A., Grieder, T., and van der Kooy, D. (2009). GABA A receptors mediate the opposing roles of dopamine and the tegmental pedunculopontine nucleus in the motivational effects of ethanol. European Journal of Neuroscience 29, 1235–1244.

37. Ting-A-Kee, R., Vargas-Perez, H., Mabey, J.K., Shin, S.I., Steffensen, S.C., and van der Kooy, D. (2013). Ventral tegmental area GABA neurons and opiate motivation. Psychopharmacology 227, 697–709.

38. Tyzio, R., Nardou, R., Ferrari, D.C., Tsintsadze, T., Shahrokhi, A., Eftekhari, S., Khalilov, I., Tsintsadze, V., Brouchoud, C., Chazal, G., et al. (2014). Oxytocin-Mediated GABA Inhibition During Delivery Attenuates Autism Pathogenesis in Rodent Offspring. Science 343, 675–679.

39. Vargas-Perez, H., Ting-A-Kee, R., Walton, C.H., Hansen, D.M., Razavi, R., Clarke, L., Bufalino, M.R., Allison, D.W., Steffensen, S.C., and van der Kooy, D. (2009). Ventral Tegmental Area BDNF Induces an Opiate-Dependent-Like Reward State in Naive Rats. Science 324, 1732–1734.

40. Vargas-Perez, H., Bahi, A., Bufalino, M.R., Ting-A-Kee, R., Maal-Bared, G., Lam, J., Fahmy, A., Clarke, L., Blanchard, J.K., Larsen, B.R., et al. (2014). BDNF Signaling in the VTA Links the Drug-Dependent State to Drug Withdrawal Aversions. Journal of Neuroscience 34, 7899–7909.

41. Volman, S.F., Lammel, S., Margolis, E.B., Kim, Y., Richard, J.M., Roitman, M.F., and Lobo, M.K. (2013). New Insights into the Specificity and Plasticity of Reward and Aversion Encoding in the Mesolimbic System. Journal of Neuroscience 33, 17569–17576.

42. Wellershaus, K., Degen, J., Deuchars, J., Theis, M., Charollais, A., Caille, D., Gauthier, B., Janssen-Bienhold, U., Sonntag, S., Herrera, P., et al. (2008). A new conditional mouse mutant reveals specific expression and functions of connexin36 in neurons and pancreatic beta-cells. Experimental Cell Research 314, 997–1012.

